# A highly active phosphate-insensitive phosphatase is widely distributed in nature

**DOI:** 10.1101/2021.08.27.457942

**Authors:** Ian D.E.A. Lidbury, David J. Scanlan, Andrew R. J. Murphy, Joseph A. Christie-Oleza, Maria M. Aguilo-Ferretjans, Andrew Hitchcock, Tim Daniell

**Affiliations:** Department of Animal and Plant Sciences, University of Sheffield, Sheffield, UK; School of Life Sciences, University of Warwick, Gibbet Hill Road, Coventry, UK; University of the Balearic Islands, Palma, Spain; Department of Molecular Biology and Biotechnology, University of Sheffield, Sheffield, UK

## Abstract

The regeneration of bioavailable phosphate from immobilised organophosphorus represents a key process in the global phosphorus cycle and is facilitated by enzymes known as phosphatases. Most bacteria possess at least one of three major phosphatases, known as PhoA, PhoX and PhoD, whose activity is optimal under alkaline conditions. The production and activity of these three phosphatase families is negatively regulated by phosphate availability and thus these enzymes play a major role in scavenging phosphorus only during times of phosphate scarcity. Here, we reveal a previously overlooked phosphate-insensitive phosphatase, PafA, prevalent in *Bacteroidetes*, which is highly abundant in nature and represents a major route for the remineralisation of phosphate in the environment. Using *Flavobacterium johnsoniae* as the model, we reveal PafA is highly active towards phosphomonoesters. Unlike other major phosphatases, PafA is fully functional in the presence of its metabolic product, phosphate, and is essential for growth on phosphorylated carbohydrates as a sole carbon source. PafA, which is constitutively produced under all growth conditions tested, rapidly remineralises phosphomonoesters producing significant quantities of bioavailable phosphate that can cross feed into neighbouring cells. *pafA* is both abundant and highly expressed in the global ocean and abundant in plant rhizospheres, highlighting a new and important enzyme in the global phosphorus cycle with applied implications for agriculture as well as biogeochemical cycling. We speculate PafA expands the metabolic niche of *Bacteroidetes* by enabling utilisation of abundant organophosphorus substrates in the presence of excess phosphate, when other microbes are rendered incapable.

**Significance statement:** Phosphorus is an essential element for all life on Earth. Global primary production, and thus the ability for oceans and soils to drawdown atmospheric carbon dioxide, is in part controlled by the availability of inorganic phosphate. Likewise, global food production is also reliant on adequate supplies of phosphorus to both plants and animals. A major fraction of the total phosphorus pool exists as organic phosphorus, which requires mineralisation to phosphate prior to incorporation into cellular biomolecules. This important process is performed by enzymes known as phosphatases. Here, we reveal that the unique bacterial phosphatase, PafA, is a key player in the global phosphorus cycle and presents a major route for the regeneration of bioavailable phosphate required for both primary and secondary production.

## Background

Both terrestrial and aquatic biological production is regulated by the availability of phosphorus (P) with consequences for global food production, biodiversity, and drawdown of atmospheric CO2^1, 2^. Thus, the global P cycle plays a vital role in sustaining human existence both presently and going into the future. P limitation is predicted to constrain the stimulation of land plant biomass in response to elevated atmospheric CO2^2, 3^. Likewise, most of the global ocean is P deplete, which can either lead to growth limiting or Pi stress conditions, reducing phytoplankton production^4^. In terrestrial and marine biomes, a large fraction of the total P pool consists of organic compounds, such as phosphomono-, phosphodi-, and phosphotri- esters, phosphonates, and phytic acid^5–7^. Remineralisation of organic P into inorganic phosphate (Pi), by either a primary producer or associated microorganisms, enhances production through alleviation of P starvation^4, 8, 9^. However, the environmental distribution of organic P mineralising enzymes and the relative contribution of distinct microbial taxa towards this process is limited^4, 10^. This reduces our ability to predict the influence of anthropogenic-driven global change on marine and soil P cycling, its interaction with the global carbon (C) cycle, and the development of sustainable agricultural tools promoting more efficient crop and animal production.

Bacteria, like all organisms on Earth, require P for survival, growth and reproduction. Their preferred source of exogenous P is Pi. However, in the environment Pi is frequently found at very low or growth limiting concentrations. Common to bacteria is the ability to overcome environmental Pi scarcity through expression of various genes encoding Pi-stress response proteins^11^. This includes the Pi-dependent production of periplasmic or outer membrane bound enzymes called phosphatases, which cleave off the Pi moiety from various organic P compounds. Typically, phosphatases target either organic phosphomonoesters, such as sugar phosphates, or phosphodiesters, such as DNA and phospholipids. Phosphatases can be separated into different classes based on their substrate range (promiscuous or specific), substrate preference (phosphomono-, phosphodi-, and phosphotri-esterases) and their pH optimum (acid or alkaline)^10–13^. The most common class of bacterial phosphatases are promiscuous alkaline phosphomonoesterases (PMEs), which can be further separated into three major families, PhoA, PhoD and PhoX^14–16^. In addition, there is a growing body of evidence that these alkaline PMEs are also active against phosphodi- and phosphotri-esters broadening the role of these enzymes^17, 18^. The unifying function of these enzymes is the production of Pi during times of Pi-depletion and consequently their regulation and enzyme activity are inhibited by ambient concentrations of inorganic Pi^19–22^. A unique but understudied fourth class of promiscuous bacterial alkaline PME also exists^23^ and was recently shown to be highly prevalent in the genomes of *Bacteroidetes*^24^. Unlike PhoA, PhoD and PhoX, this enzyme, referred to as PafA, is not repressed by Pi at either the regulatory or enzyme activity level^18, 23^. Therefore, whilst the metabolic role of PafA is unknown, the regulatory and biochemical data suggests its function is greater than scavenging Pi during times of Pi-limitation.

In marine, ocean, and soil microbiomes, members of the phylum *Bacteroidetes* are major degraders of plant and algal glycans occupying a functional niche focused on the degradation of high molecular weight organic polymers^25–27^. Recently, plant associated *Bacteroidetes* have been shown to play a major role in supressing plant disease and there is a growing interest in their ability to augment plant nutrition^24, 28–30^. A defining genomic signature of *Bacteroidetes* is the possession of specialised outer membrane transporters, commonly referred to as SusCD (archetypal transporter is the C and D components of the **<u>S</u>**tarch **<u>U</u>**tilisation **<u>S</u>**ystem), that facilitate the uptake of large polymers as nutrition^31^. This is coincident with the apparent lack of ATP-binding cassette (ABC) transporters required for the active transport of smaller molecules. Therefore, in addition to specialising in the degradation of HMW polymers, *Bacteroidetes* must possess fundamentally different molecular mechanisms to capture nutrients in comparison to other bacterial taxa. Members of the Bacteroidetes phylum, predominantly *Flavobacteraceae* and *Sphingobacteraceae*, are heavily enriched in the plant microbiome relative to their abundance in surrounding bulk soil communities. In the rhizosphere and root endosphere, *Bacteroidetes* can represent over half of the total microbial community. Thus, this phylum must be competitive for various growth limiting nutrients such as C, N and P despite an apparent lack of transport systems required for nutrient acquisition.

Recently, we discovered plant-associated *Flavobacterium* spp. possess remarkable potential to mobilise organic P^24^. This included the synthesis of numerous phosphomonoesterases (PMEs) and phosphodiesterases (PDEs) and the induction of novel P-regulated gene clusters, some of which harboured SusCD-like transporters, termed <u>P</u>hosphate <u>U</u>tilisation <u>S</u>ystem (PusCD). In addition, numerous *Flavobacterium* spp, especially plant-associated strains, lacked the high affinity phosphate ABC transporter, strengthening the hypothesis that this phylum have divergent mechanisms for nutrient acquisition. Another key characteristic of soil *Bacteroidetes* was the constitutive production of PME activity even in the presence of excess exogenous Pi. Through protein fractionation using *Flavobacterium johnsoniae* as the model, PafA was identified as the likely candidate for this unusual PME activity^24^, which agrees with previous enzyme kinetic studies on this PME^23^.

Here, by utilising bacterial genetics we aimed to identify the contribution of various predicted PMEs towards PME activity using *F. johnsoniae* as the model. We also tested the hypothesis that the Pi-irrepressible PME, PafA, has a primary role other than Pi scavenging during growth limiting P conditions. We discovered PafA identified in soil *Bacteroidetes* is highly active and in the absence of an ABC transport system is essential for growth on sugar phosphate phosphomonesters as a sole C and energy source.

## Results

### *Flavobacterium* PafA is a highly active phosphomonoesterase

The model bacterium *F. johnsoniae* DSM2064 (DSM2064) produces four alkaline PMEs (encoded by *fjoh_0023*, *fjoh_3187*, *fjoh_3249*, *fjoh_2478*) that are related to previously characterised PMEs (Fig. 1a)^24^. The first (encoded by *fjoh_2478*) is a lipoprotein distantly related to PhoX, which was abundantly secreted in response to Pi-depletion. Two periplasmic PMEs (encoded by *fjoh_3187* and *fjoh_3249*) were related to PhoA. The last, (encoded by *fjoh_0023*) is a Pi-irrepressible phosphatase PafA^23^. PafA is an unusual PME that has the pfam domain 01663, typically associated with phosphodiesterase activity, and does indeed mineralise both phosphomono- and phosphodiesters^18^. Indeed, PafA is most closely related to PhoD (Fig. 1a), an enzyme that is primarily a phosphodiesterase^20, 32^, whilst the Pi- irrepressible phosphatase identified in *Zymomonas mobilis*, labelled as a PhoD^33^, is also most closely related to *Bacteroidetes* PafA (Fig.1a). Two of these *Flavobacterium* PMEs, encoded by *fjoh_0023* and *fjoh_3187*, were enriched in protein fractions expressing high PME activity^24^.

**Figure 1.**
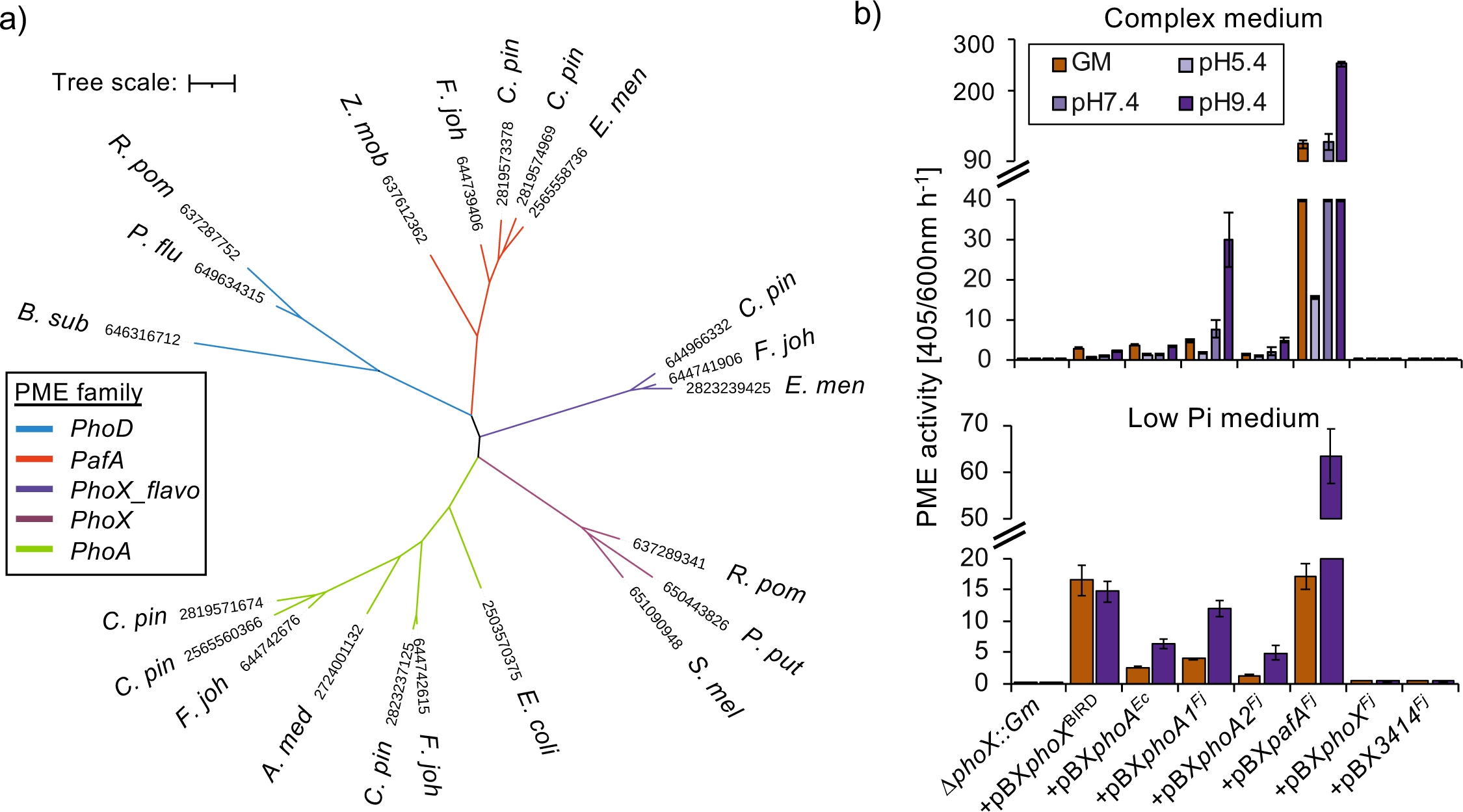
(a) Phylogeny and phosphomonoesterase (PME) activity for members of the major bacterial alkaline phosphatase families. Un-rooted phylogenetic tree comparing PhoD, PhoX, PhoA with PafA homologs. Tree topology and branch lengths were calculated by maximum likelihood using the Blosum62+F+G4 model of evolution for amino acid sequences based on 875 sites (595 parsimony-informative) in lQ-TREE software. A consensus tree was generated using 1000 bootstraps. Abbreviations: *F. joh*, *Flavobacterium johnsoniae*; *C. pin*, *Chitinophaga pinensis*; *E. men Elizabethkingia meningoseptica*; *R. pom*, *Ruegeria pomeroyi*; *B sub*, *Bacillus subtilis*; *P. flu*, *Pseudomonas fluorescens*, *P. put*, *Pseudomonas putida*, *Z. mob*, *Zymomonas mobilis*; *S. mel, Sinorhizobium meliloti*; *A. med*, *Alteromonas mediterranea* **(b)** PME activity for a *Pseudomonas putida* Δ*phoX::Gm* <u>mutant complemented with</u> various PMEs from *P. putida* BIRD (+pBX*phoX*^BIRD^), *Escherichia coli* (+pBX*phoA^Ec^*), and *F. johnsoniae* (e.g. +pBX*pafA^Fj^*) was recorded in cell cultures grown overnight (n=3) in complex medium (upper panel) or minimal medium established phosphate-deplete (Low Pi) growth conditions (lower panel). PME activity was obtained through addition of the artificial substrate *para-*nitrophenyl phosphate (10mM) under four conditions: the original growth medium (GM) or by resuspending cells in a buffer adjusted to three pH values. Values presented are the mean of biological triplicates and error bars denote standard deviation.

To investigate the relative activity of these *Flavobacterium* PMEs and directly compare them with the classical PhoX (*Pseudomonas putida*) and PhoA (*Escherichia coli*), we cloned their respective genes into a heterologous host (*Pseudomonas putida* BIRD-1), lacking a functional PME (Δ*phoX::Gm-BIRD-1*) to remove its innate PME activity^34^. All PMEs were under the control of the native *phoX* promoter encoded on the broad host range plasmid pBBR1MCS-km. This promoter was previously shown to be regulated in a PhoBR-dependent manner^19^, but PhoX peptides were still detected under Pi-replete conditions, suggesting some residual level of *phoX* expression. The various complemented Δ*phoX*^BIRD^*::Gm* strains were grown overnight in either a complex medium (Fig. 1b) or a minimal medium containing a growth limiting concentration of Pi (50 µM) (Fig. 1c). PME assays were performed using culture suspensions or by resuspending cells in a buffer at three pH conditions (5.4, 7.4, 9.4). Cells producing well-known PMEs, *i.e.* the native *P. putida* BIRD-1 PhoX (+pBX*phoX*^BIRD^) or *E. coli* PhoA (+pBX*phoA^Ec^*), both restored PME activity in the *Pseudomonas* mutant (Fig. 1b). Cells producing the *Flavobacterium* PhoA homologs, encoded by *fjoh_3249* (+pBX*phoA1^Fj^*) or *fjoh_3187* (+pBX*phoA1^Fj^*) also restored PME activity, confirming their function as PMEs. Notably, cells producing PafA (+pBX*pafA^Fj^*), encoded by *fjoh_0023*, had significantly greater activity when grown in either growth medium, especially the complex medium (Fig. 1B). Therefore even under Pi-replete growth conditions, when synthesis of PafA is low this enzyme is capable of rapidly mineralising organophosphorus (Fig. 1b). Indeed, both liquid cultures and surface-attached colonies demonstrated phosphate-dependent inhibition of *phoX*^BIRD^ or *phoA^Ec^* PME activity (Fig. 1b & Fig. S1). Cells complemented with *fjoh_*2478 (+pBX*phoX^Fj^*) failed to produce any detectable activity, which we attribute to differences between *Pseudomonas* (Type II secretion system) and *Flavobacterium* (Type IV secretion system) lipoprotein export mechanisms, leading to a non-functional polypeptide.

### PafA significantly contributes towards PME activity in *F. johnsoniae* DSM2064

Next, we examined the contribution of these PMEs towards activity in the original strain DSM2064. As expected^24^, wild type cells produced both constitutive and elevated inducible PME activity (Fig. 2). Interestingly, mutation of *pafA* (*fjoh_0023*) significantly reduced constitutive and inducible PME activity, revealing this gene plays a major role under both growth conditions. We hypothesised that a gene (*fjoh_0074*) encoding a constitutively produced lipoprotein, predicted to be a member of the endonuclease/exonuclease/phosphatase superfamily (InterPro#; PR005135), may make a minor contribution to PME activity under Pi-replete growth conditions. However, mutation of this gene had no apparent effect on the observable PME activity in DSM2064. Mutation of *fjoh_2478* (Δ*phoX*), encoding the distinct PhoX-like lipoprotein (Fig. 1a), reduced inducible PME activity in DSM2064. Likewise, a double mutant (Δ*fjoh_3249:*Δ*fjoh3187*) defective for the two *phoA*-like (ΔA1: ΔA2) genes also reduced inducible PME activity. Through generation of a quadruple knockout PME mutant, Δ*fjoh_0074:*Δ*fjoh_2478:*Δ*fjoh_3249:*Δ*fjoh3187* (Δquad), and a quintuple knockout PME mutant, Δ*fjoh_0074:*Δ*fjoh_2478:*Δ*fjoh_3249:*Δ*fjoh3187:*Δ*pafA* (Δquad: ΔpafA), we confirmed that 1) *pafA* contributes to >95% of the constitutive and approximately half the inducible PME activity and 2) *fjoh_2478* (*phoX*), *fjoh_3249* (*phoA1*) and *fjoh_3187* (*phoA2*) are responsible for the additional inducible PME activity expressed in response to Pi-depletion. Complementation of the quintuple PME mutant with its native *pafA* (+pY:pafA) duly restored PME activity to that comparable with the quadruple PME mutant, confirming the above results. Finally, mutation of *pafA* (*fjoh_0023*) in the Δ*phoX* background, creating a double mutant strain (Δ*fjoh_2478:*Δ*fjoh_0023*), further confirmed a role for *fjoh_2478* in inducible PME activity (Abs405/600 nm h^-1^: Δ*pafA* = 21±/-2.8; Δ*pafA:*Δ*phoX* = 9.6±1.2). Together, DSM2064 possess four PMEs that contribute towards PME activity against the artificial substrate *p*NPP. There is also evidence from the quintuple mutant that another unidentified PME has a very minor contribution towards PME activity.

**Figure 2.**
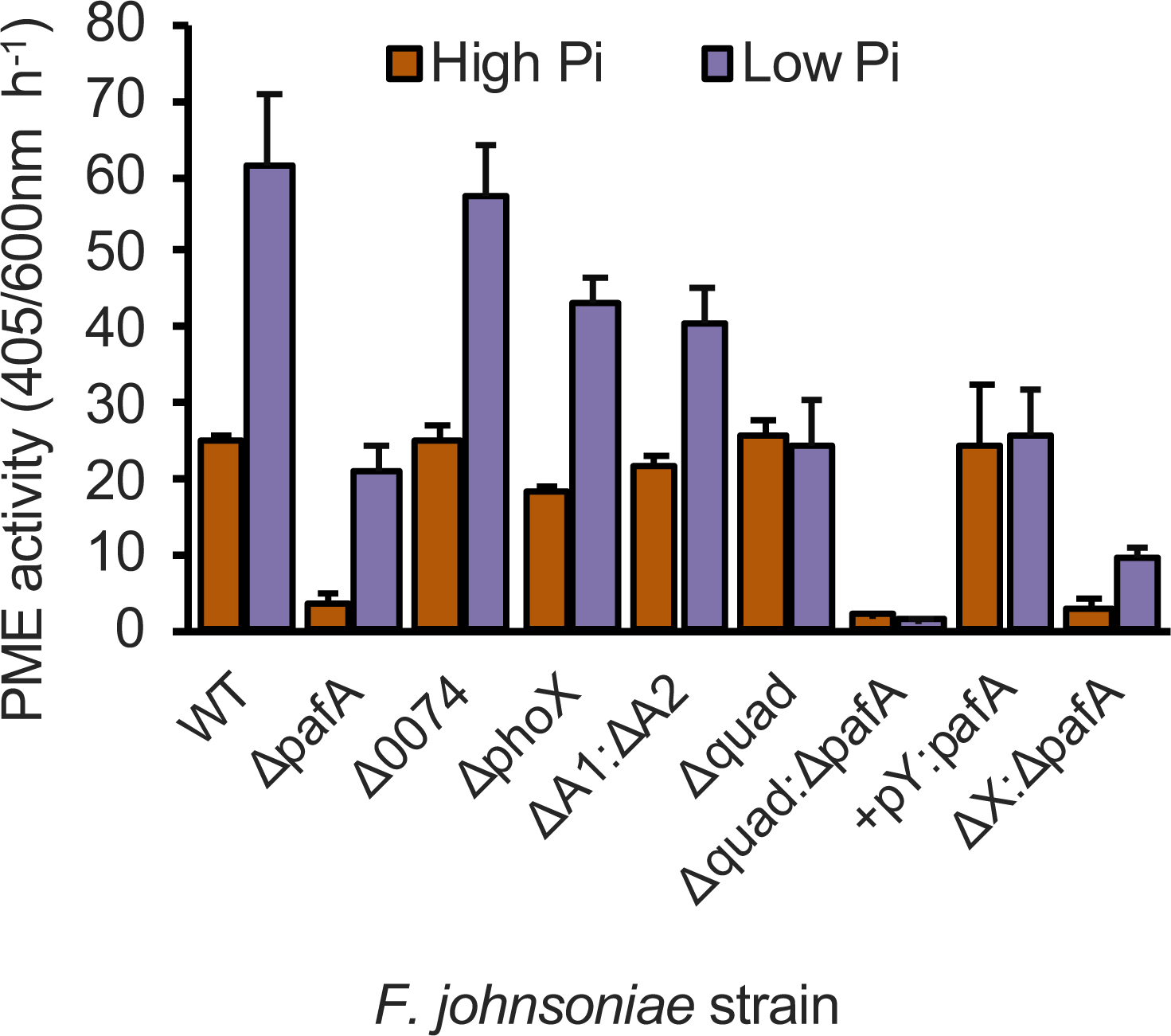
Phosphomonoesterase (PME) activity in various *F. johnsoniae* genotypes. PME activity was recorded in cell cultures (n=3) grown overnight under phosphate-replete (High Pi) or phosphate-deplete (Low Pi) growth conditions. Activity was obtained through addition of the artificial substrate *para*-nitrophenol phosphate (10mM). Values presented are the mean of biological triplicates and error bars denote standard deviation. Abbreviations: WT, wild type; A1:A2, double *phoA* mutant; quad, quadruple mutant; quad:pafA, quintuple mutant; +pY:pafA, quintuple mutant complemented with *pafA*; X:pafA, *phoX pafA* double mutant.

### PafA enables utilisation of organic P substrates as both a P and a C source

To determine the functional role of PafA in organic P mineralisation, we screened the parental wild type and selected PME mutant strains of DSM2064 (Fig. 3) on a range of organic P substrates as the sole P source (300 μM). Wild type growth on organic P substrates was comparable to growth on Pi whilst growth of the quintuple PME mutant was severely inhibited (Fig. 3a). In agreement with the very low but observable level of PME activity in the quintuple PME mutant, a small but detectable amount of growth did occur relative to the no P control. The *pafA* complemented quintuple PME mutant (+pY:*pafA*) fully restored the wild type phenotype (Fig. 3a), indicating PafA alone can facilitate utilisation of organic P as a sole P source despite its relatively low level of expression^24^. A double mutant, defective for *pafA* and *phoX*^2064^ (Δ*fjoh_2478:*Δ*fjoh_0023*) had significantly reduced and variable rates when organic P was the substrate indicating the two *phoA*-like homologs play a minor role in growth on these organic P substrates.

**Figure 3.**
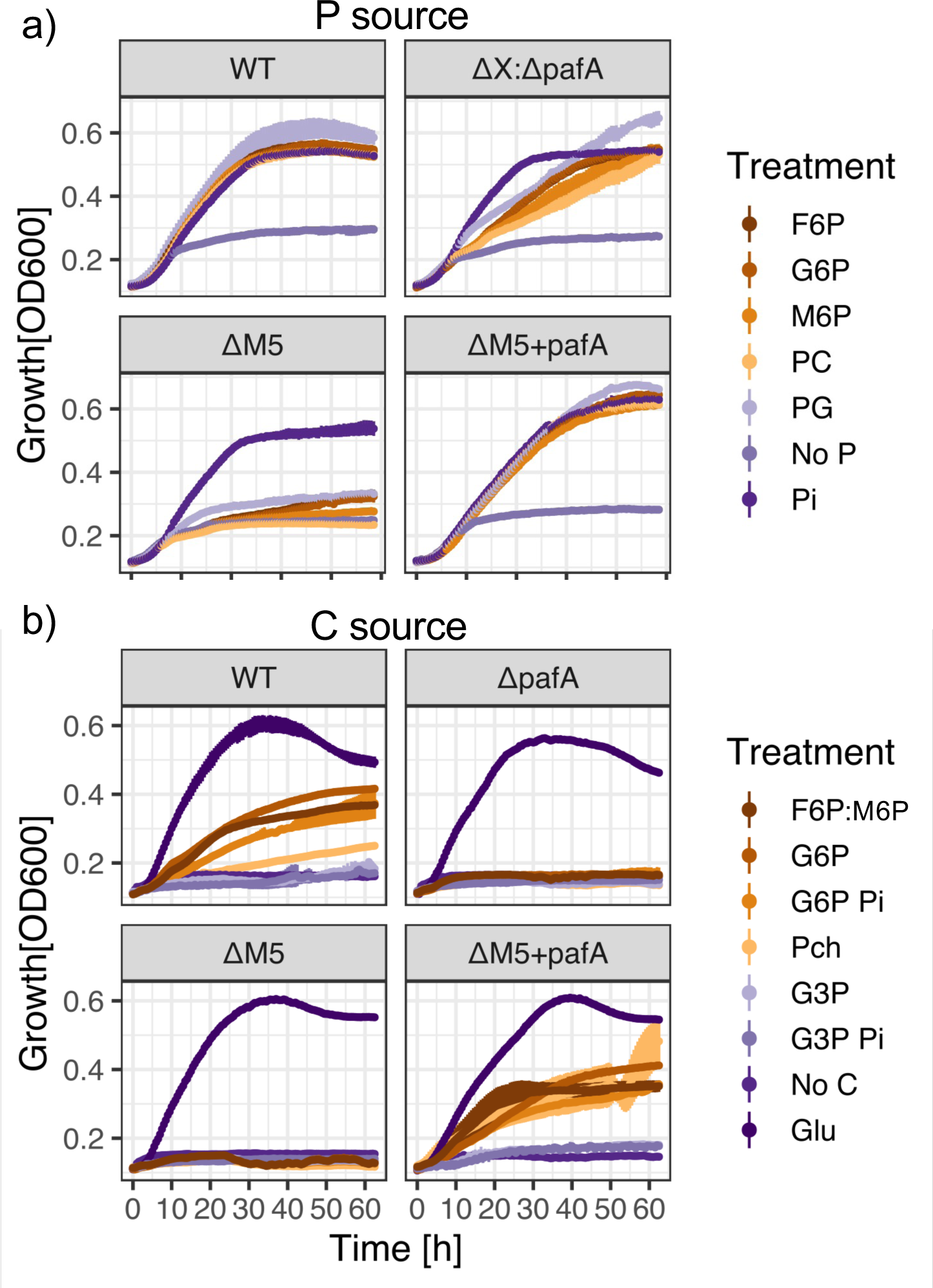
Growth of *F. johnsoniae* DSM2064 on various organic P substrates as a sole P and C source. (a) The wild type (WT), Δ*fjoh_2478:*Δ*fjoh_0023* (ΔX:ΔpafA) double mutant, quintuple mutant (M5) and complemented quintuple mutant with *pafA* (M5+pafA) were grown (n=3) on various P substrates (200 µM) in addition to a no P control. (b) The WT, *fjoh_0023* (ΔpafA) single mutant, M5 and M5+pafA were also grown (n=3) on various P substrates (3 mM) as the sole C source. Results presented are mean values and error bars denote standard deviation. Abbreviations: F6P, fructose 6-phosphate; G6P, glucose 6- phosphate; M6P, mannose 6-phosphate; PC, phosphorylcholine; PG, phosphoglycerol; Pi phosphate; No P, negative control.

Next, we investigated whether PafA provides a novel mechanism for growth on organic P substrates as a sole C, P, and energy source. To do this, 5 mM of various organic P substrates were supplied as the sole C and P source (Fig. 3b). In addition, glycerol 3-phosphate and glucose 6-phosphate were provided as a sole C source but also supplemented with Pi (1mM) to inhibit expression of inducible PMEs. The wild type efficiently utilised phosphorylated carbohydrates (mannose 6-phosphate, fructose 6-phosphate and glucose 6- phosphate) as a sole C and P source, grew slowly on phosphocholine (PC), but could not utilise glycerol 3-phosphate. The addition of exogenous Pi (inhibiting expression and activity of inducible PMEs) did not affect growth, suggesting PafA was the major enzyme required for this phenotype. Indeed, the *pafA* single mutant and quintuple PME mutant failed to grow on these organic substrates as a sole C source and complementation of the quintuple PME mutant with *pafA* fully restored the wild type phenotype, confirming the essential role of *pafA* in the utilisation of organic P substrates as a sole C and energy source.

### Organic P mineralisation by *Flavobacterium* facilitates cross feeding of bioavailable P

To determine the ecological consequences of Pi-independent organic P mineralisation in DSM2064, we grew the quintuple PME mutant, impaired in its ability to utilise various organic P substrates (Fig. 3), in co-culture with the parental wild type using various organic P substrates as the sole P source (300 μM). To achieve a semi-continuous growth pattern, after 24-48 h growth each culture line was twice inoculated (2% v/v) into fresh medium containing the same organic P substrate. The control treatment, using Pi as a sole P source resulted in comparable growth between the quintuple PME mutant and wild type (Fig. 4a) and thus a similar ratio between the two strains (Fig. 4b). When phosphorylated carbohydrates, such as fructose 6-phosphate or glucose 6- phosphate were supplied as the sole P source, the mutant also grew at a comparable rate to the wild type, with strains maintaining a 1:1 ratio. However, when phosphocholine or the phosphodiester-containing lipid phosphatidylinositol were the sole P source the mutant was significantly outcompeted by the wild type (PI =31.4±9-fold, PC =15.9±6-fold). This would suggest that for *Flavobacterium*, different organophosphorus compounds may be mineralised at distinct locations, i.e. extracellular or periplasmic, resulting in inefficient versus efficient acquisition, respectively.

**Figure 4.**
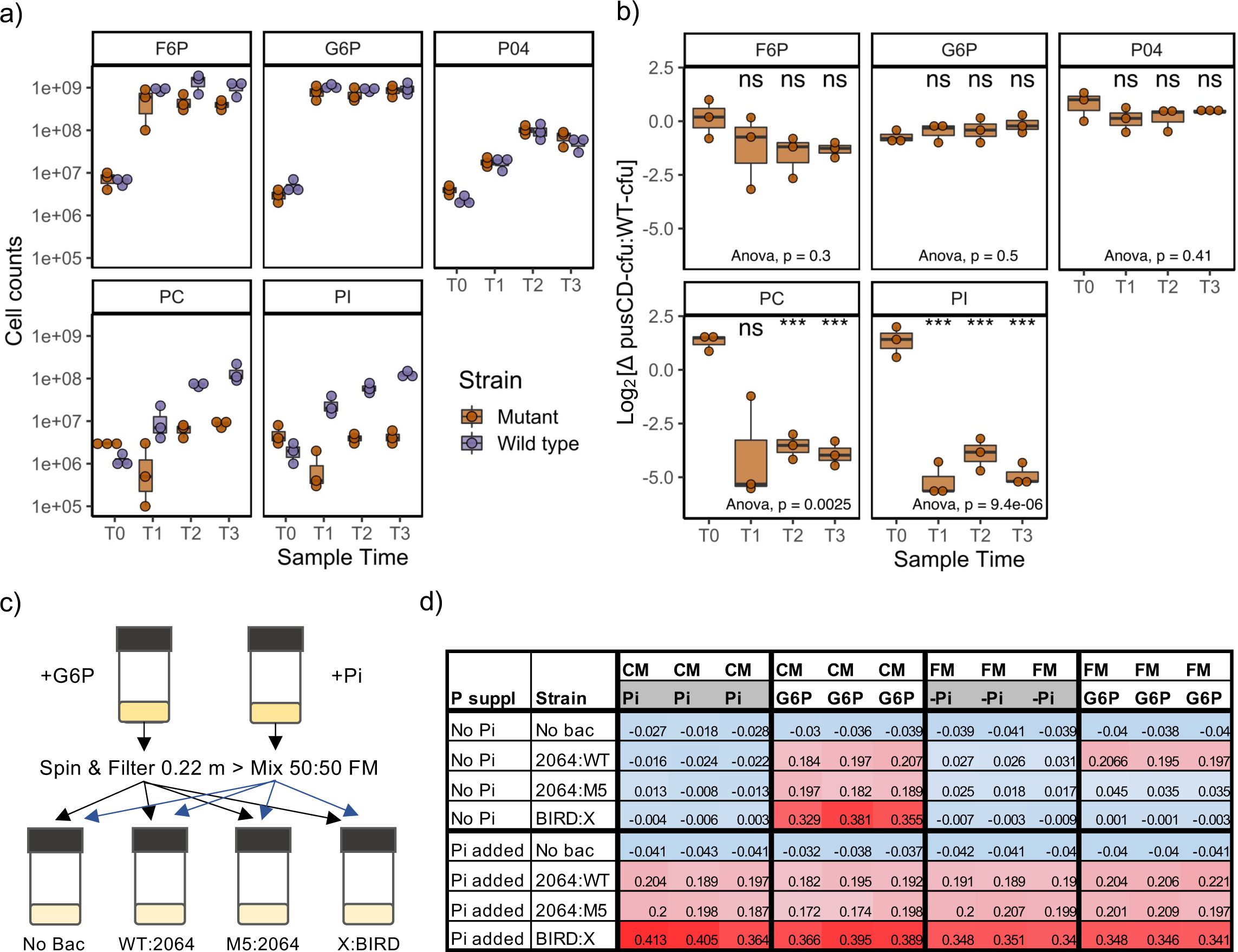
Co-culture growth experiments using organic P growth substrates as the sole P source. *F. johnsoniae* wild type and quintuple mutant were grown in competition for the organic P substrates fructose 6-phsphate (F6P), glucose 6-phopshate (G6P), phosphorylcholine (PC) and the phospholipid, phosphatidylinositol, and inorganic phosphate (PO4) as the sole P source. **(a)** Enumeration of the growth of each strain was calculated by obtaining colony forming units and **(b)** the fitness of the mutant relative to the wild type was determined by plotting the difference in counts. Cultures were grown in triplicate (individual data points displayed) and box whiskers represent the mean, first and third quartiles and upper and lower interquartile ranges. **(c)** Schematic for the generation of conditioned medium (CM) and screening for the accumulation of mineralised phosphate in the growth medium including removal of wild type cells after initial growth and exhaustion of carbon and energy. **(d)** Growth of either wild type *F. johnsoniae* (WT:2064), the quintuple PME mutant (M5:2064) or *phoX:Gm-BIRD-1* (X:BIRD) on CM or fresh medium containing either organic phosphorus or phosphate. Values represent OD600 of cultures and the shading is a visual representation of growth; blue: red scale equals low to high growth.

Next, to investigate whether growth on phosphorylated carbohydrates as a sole C and energy source would result in the accumulation of excess Pi in the medium, we grew the wild type in minimal medium supplemented with glucose 6-phosphate (2mM) and glucose (5mM). This achieved a low C:P ratio (21:1), well below cell stoichiometric requirements (106:1), that should facilitate accumulation of Pi in response to continual Pi-independent mineralisation by PafA. We also established a control treatment containing glucose (10 mM) and Pi (25 μM) that would create Pi-deplete growth conditions (C:P, 2400:1) and thus cause complete consumption of exogenous Pi. After overnight growth, cells were removed via filtration and the conditioned medium (CM) was mixed (50:50% v/v) with fresh medium (FM) containing 10 mM glucose and no added P source (Fig. 4c). Each cell-free growth medium was then either inoculated with the wild type, the quintuple PME mutant or the Δ*phoX::Gm-BIRD-1* mutant. Control treatments using only FM supplemented with either glucose 6-phosphate or Pi and an additional positive control with Pi supplemented to all growth condition-strain combinations were established. Cultures with no added bacterial inoculant had no growth, confirming cells were successfully removed from the initial conditioned medium (Fig. 4d). All positive control cultures supplemented with Pi all grew for both FM and CM treatments (bottom half Fig. 4d). For FM cultures, all strains failed to grow in the absence of Pi and only wild type DSM2064 grew in the presence of glucose 6-phosphate whilst the quintuple PME mutant or the *P. putida* mutant did not. No growth was observed for any strain using CM originally containing 25 μM Pi whilst all three strains grew in CM originally containing 2mM glucose 6-phosphate. Together, these data clearly demonstrate mineralisation of organic P independently of cellular P requirements and production of bioavailable Pi for other organisms.

### PafA facilitates the rapid mineralisation of organic P substrates

Finally, to quantify the contribution of PafA and other PMEs towards the mineralisation of organic P, the concentration of exogenous Pi in the culture supernatant was quantified during growth on organic P. DSM2064 wild type and the various PME mutants (Δ*pafA*, Δ*pafA:*Δ*phoX*, or quintuple PME) were individually grown on glucose 5 mM supplemented with either 2 mM glycerol phosphate and phosphocholine (50:50 mix); or 2 mM phosphorylated carbohydrates (50:50 mix G6P:F6P) as well as a Pi control (500 μM). After 25 h, cultures were further supplemented with another 5 mM glucose. All four DSM2064 genotypes grew at comparable growth rates, although the quintuple PME mutant showed impaired growth on organic P relative to the wild type, as expected (Fig. 5). We attribute this growth to the higher concentration of organic P that likely resulted in a higher trace contamination of Pi. Indeed, detectable concentrations of Pi (14-24 μM) were detected at T0 in cultures supplemented with organic P only. After 5 h, all cultures showed minimal growth. However, wild type cultures grown in the presence of PG:PC or F6P:G6P had already accumulated 384±53 μM and 564±17 μM of exogenous Pi, respectively by this time (Fig. 5). All three PME mutant strains lacking a functional PafA showed no significant release of remineralised Pi. After 25 h growth, wild type cultures accumulated >1 mM Pi, whilst no Pi was detected in the quintuple PME mutant. The Δ*pafA* mutant possessing a functional PhoX did produce significantly more Pi in the presence of F6P:G6P compared to PG:PC, unlike the Δ*pafA*:Δ*phoX* double mutant that lacks a functional PhoX. This observation may explain why phosphorylated carbohydrates cross feed into non-utilisers, unlike phosphocholine or phosphatidylinositol, that appear to be more efficiently utilised (Fig. 4). In summary, these data reveal PafA plays a major role in the mineralisation of organic P and presents a rapid route for production of exogenous bioavailable Pi.

**Figure 5.**
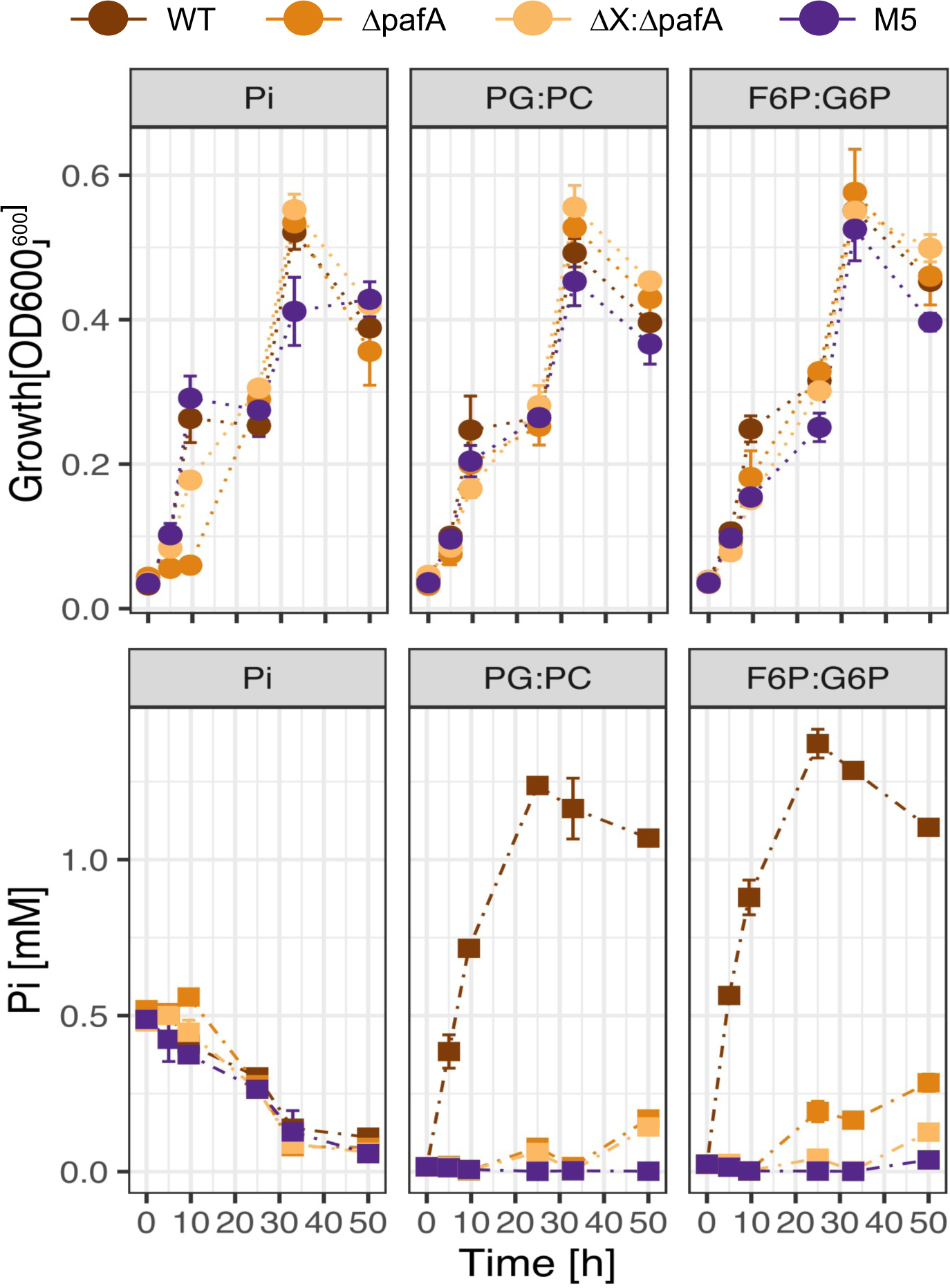
Organic P mineralisation and Pi export rates in various *F. johnsoniae* DSM2064 genotypes. Wild type and three PME mutants were grown in the presence of glucose (5mM) and an organic P substrate mix (2 mM). After 25 h, an additional 5 mM glucose was supplemented to all cultures. Growth (upper panel) and Pi accumulation (lower panel) were quantified over time. Data points represents the mean of duplicate cultures sampled in duplicate (n=4) and error bars denote the standard deviation. Abbreviations: Pi, phosphate; PG:PC, phosphoglycerol-phosphocholine mix; F6P:G6P, fructose 6-phosphate-glucose 6- phosphate mix; M5, quintuple PME mutant.

### PafA diversity and occurrence in *Bacteroidetes* and non-*Bacteroidetes* bacteria

Whilst the majority of inducible PMEs are predominantly found in the genomes of plant-associated *Bacteroidetes*, the occurrence of PafA in bacteria related to this phylum is more widespread^24^. To further investigate the environmental distribution of PafA, we scrutinised various marine (TARA Oceans), rhizosphere (Oil Seed Rape and Wheat) and gut (rumen and human) metagenomes (total n = 357). We combined the retrieved environmental sequences with the PafA sequences previously identified in our *Bacteroidetes* isolate genome bank (n = 468)^24^. Only sequences possessing the key residues^18, 24^ required for PME activity were retained (Environmental n=1314; isolate n=423). Phylogenetic analysis revealed the existence of several polyphyletic subclades, as evidenced by the occurrence of two distinct PafA homologs found in the model soil bacterium *Chitinophaga pinensis* (Cp1 and Cp2 in Fig. 6a), that were partially resolved by their environmental origin (isolate or metagenome). To confirm the function of the *pafA* homologs within these branches, we cloned both *C. pinensis* genes encoding Cp1 and Cp2 into the same expression vector and heterologous host system. Both were functional Pi-insensitive PMEs (Fig S2) and could mineralise naturally-occurring organic P molecules (Fig. 6b).

**Figure 6.**
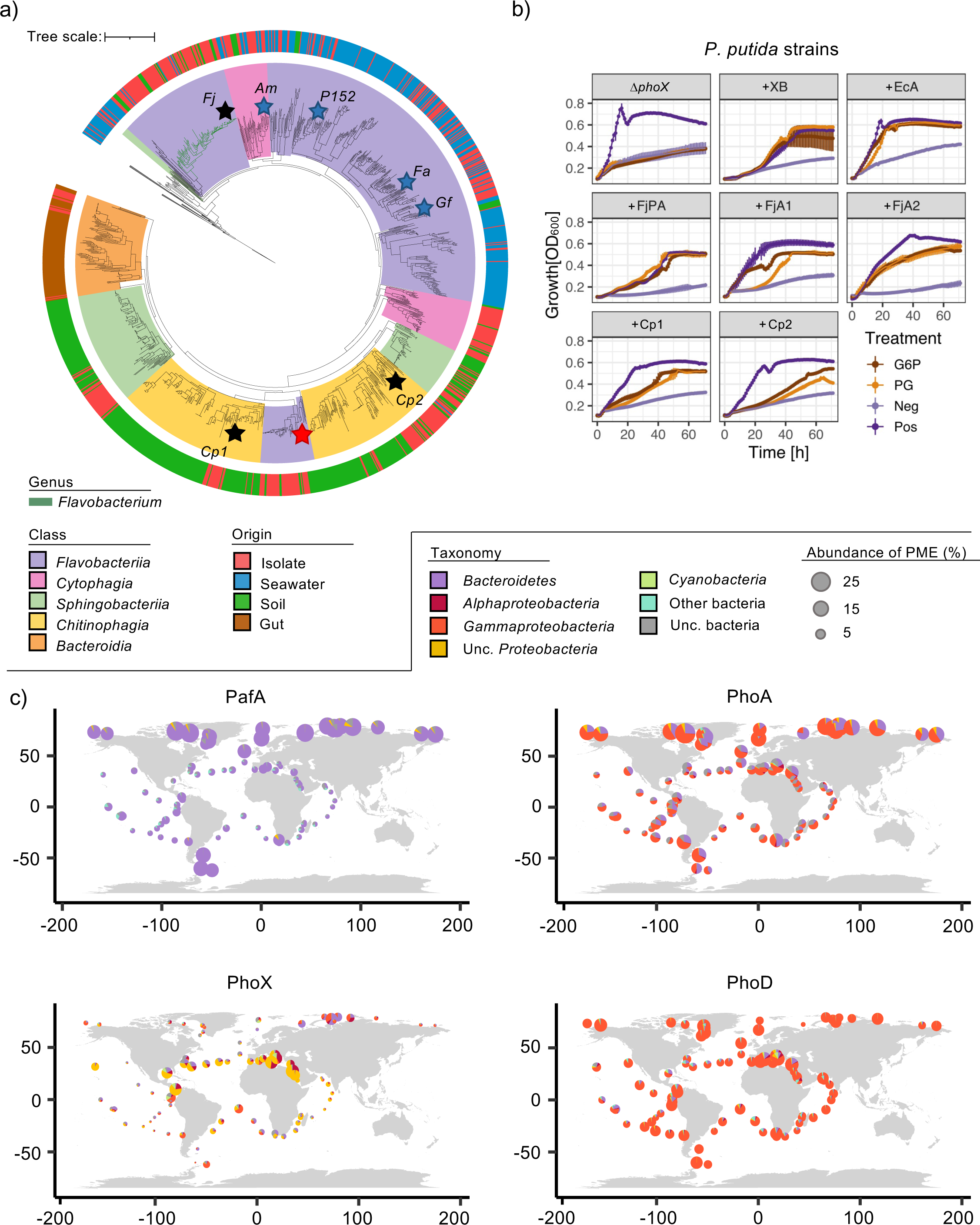
Environmental diversity of PafA and other alkaline PMEs in soil, gut and ocean microbiomes. (a) PafA diversity in genome-sequenced isolates and environmental metagenomes (outer ring denotes origin). PafA sequences related to each class of Bacteroidetes are coloured. Branches corresponding to the genus *Flavobacterium* are highlighted green. Black stars represent PafA homologs cloned into the heterologous host *P. putida*. The red star denotes the characterised PafA found in *Elizabethkingia meningoseptica*. **(b)** Distribution of the four major alkaline PMEs in the global ocean based on the TARA Oceans dataset. The area of each pie chart represents the normalised gene abundance, expressed as the % of bacteria possessing a given PME (see legend for scaling), at each sampling site and the contributing taxa.

Sequences related to soil/plant associated *Flavobacterium* PafA represented a small fraction of the overall PafA diversity, and few environmental sequences were retrieved from the soil/rhizosphere MGs associated with the two crop species. A large proportion of diversity belonged to marine *Flavobacteriia* (predominantly the family *Flavobacteriaceae*) including numerous sequences captured from the TARA oceans dataset. Screening four marine *Bacteroidetes* and three marine *Alphaprotoebacteria* showed that possession of *pafA* correlated with Pi-insensitive PME activity, i.e. all four marine *pafA*-encoding Bacteroidetes showed high PME activity even in high P medium whereas all three none *pafA*-encoding *Alphaproteobacteria* had their PME activity inhibited in such Pi-rich medium (Table 1). All gut related PafA sequences clustered closely with isolates related to *Bacteroidales*, including *Prevotella* spp. The previously characterised PafA from *Elizabethkingia meningoseptica* was in a cluster associated with *Chryseobacterium* sequences more closely related to *Sphingobacteraceae* sequences rather than *Flavobacteraceae* (Fig. 6a). These data suggest PafA may have undergone some level of envionrmental adaptation after its first appearance.

**Table 1.** Phosphomonoesterase (PME) activity produced by marine bacteria grown overnight (n=3) in marine Broth (complex medium, phosphate-replete), obtained through addition of the artificial substrate para-nitrophenyl phosphate (10mM).

| Strain | Phylum | Class | PME activity<br>(405/600nm h<sup>-1</sup>) |
| --- | --- | --- | --- |
| *Roseobacter denitrificans* OCh 114 | *Proteobacteria* | *Alphaproteobacteria* | 0.033 ±0.044 |
| *Ruegeria pomeroyi* DSS-3 | *Proteobacteria* | *Alphaproteobacteria* | 0.001 ±0.019 |
| *Dinoroseobacter shibae* DFL-12 | *Proteobacteria* | *Alphaproteobacteria* | 0.055 ±0.057 |
| *Algoriphagus machipongonensis* PR1<sup>†</sup> | *Bacteroidetes* | *Cytophagia* | 3.214 ±0.069 |
| *Formosa agariphila* KMM 3901<sup>†</sup> | *Bacteroidetes* | *Flavobacteriia* | 2.335 ±0.089 |
| *Polaribacter* sp. MED152<sup>†</sup> | *Bacteroidetes* | *Flavobacteriia* | 1.510 ±0.043 |
| *Gramella forsetii* KT0803<sup>†</sup> | *Bacteroidetes* | *Flavobacteriia* | 0.509 ±0.070* |

\*Poor growth and cell aggregation in culture medium
<sup>†</sup>Possession of *pafA* in the genome

### PafA and PhoA are highly prevalent in the global ocean

We next compared the abundance, expression, and associated phylogeny of the genes encoding the three major classes of bacterial alkaline PMEs (PhoA, PhoD, PhoX) and PafA in the global ocean. Unexpectedly, both *pafA* and *phoA* were more prevalent and more transcribed than *phoX* at numerous oceanic sites, particularly those in polar and temperate regions of the ocean (Fig. 6c). For example, across the Arctic Ocean and Southern Ocean, *pafA* and *phoA* transcription was over 10-fold greater than *phoX* (Fig. S3). As expected, most *pafA* sequences were related to *Bacteroidetes.* We also found an unexpected abundance of *phoD* sequences across most oceanic cites, the majority of which were related to marine *Gammaproteobacteria*. Given these unexpected results that contradict previous work^21^, we also analysed the TARA Oceans database using the latter authors bioinformatics pipeline. This revealed that BLASTP analysis dramatically underestimated the prevalence of *phoA* (Table S3) by failing to capture the full diversity of *phoA* sequences present in the global ocean (Fig. S4 and S5). *phoX* was significantly more abundant than *pafA* in the Mediterranean Sea and regions of the North Atlantic Ocean (Fig. S3), typified by very low concentrations of Pi^35^. Indeed, in these regions, all three classical PMEs were more prevalent and transcribed at significantly higher levels than *pafA* (Fig. S3). Regression analysis confirmed global *phoX* prevalence (R^2^ = 0.1804, p<0.001) and transcription (R^2^ = 0.05137, p<0.01) was negatively correlated with standing stock concentrations of Pi (Fig. S6). On the other hand, global *pafA* prevalence (R^2^ = 0.1296, p<0.001) and transcription (R^2^ = 0.05432, p<0.01) was positively correlated with Pi, demonstrating the Pi-independent functional role of this enzyme. In contrast, *phoD* and *phoX* gene abundance and expression were both negatively correlated. *phoA* prevalence did not correlate with Pi (R^2^ = 0.0028, p=0.24) though transcription was positively correlated with Pi (R^2^ = 0.06746, p<0.001). In summary, *pafA* and *Flavobacteriia*- like *phoA* sequences are diverse and widespread in nature including soil, gut and ocean microbiomes with PafA representing a major new enzyme in the global P cycle.

## Discussion

We report a unique class of alkaline phosphatase, termed PafA, is abundant in nature and enables the rapid mineralisation of various organic P substrates independently of exogenous Pi concentration. Despite *Flavobacterium* spp. possessing numerous PMEs, PafA was essential for growth on phosphorylated carbohydrates as a sole C or sole C and P source, revealing functional diversification of PafA compared to other well-known phosphatases^19–22, 36^. These findings also explain why regulation of PafA production and enzyme activity are Pi-insensitive^18, 23, 24^. Pi-insensitive mineralisation of specific reduced organophosphorus compounds (phosphonates) is widespread in the global ocean and represents a major route for regenerating the Pi required for marine primary production^37, 38^. Despite relatively low expression of *pafA* in *Flavobacterium* spp. during Pi-replete or Pi-deplete growth conditions^24^, we show this enzyme possesses superior activity towards the artificial substrate *p*NPP in comparison with PhoX or PhoA and rapidly converts various natural organic P substrates into bioavailable Pi independently of growth status. Thus, we reveal another Pi-insensitive mechanism for the rapid conversion of organic P into bio-available P, which may explain why PME activity is detected and operational in Pi-replete oceanic regions^4, 39, 40^. Significantly, *pafA* is present in the genomes of *Bacteroidetes*, who frequently associate with phytoplankton and sinking particles^41, 42^, and is expressed at levels comparable to and even exceeding other well-known phosphatases, such as *phoX*.

Traditionally, *phoX* was considered the most abundant phosphatase in the ocean14, and that iron limitation may affect this enzyme’s efficacy and limit microbial Pi mineralisation^43^. However, divergent *phoA* homologs, that do not require iron for catalytic activity^18^, are also prevalent in the global ocean^17^. Our comprehensive comparative analysis of the four major bacterial alkaline phosphatases in the global ocean revealed an unexpected distribution and diversity of these P cycling genes which significantly alters the paradigm around marine P cycling and the major players that are involved. Indeed, PafA homologs were also found in the genomes of abundant oligotrophic soil and marine bacteria related to *Acidobacteria*, *Candidatus* Lindowbacteria, V*errucomicrobia,* Gemmatimonadetes and Planctomycetes and future research should ascertain their activity and sensitivity towards Pi to improve our understanding of global P cycling. Whilst extrapolation from lab cultures can be misleading, the difference between *phoX* and *pafA* expression profiles across the global ocean further strengthens the idea that PafA has a functional role greater than that of scavenging P, i.e. a role in C-utilisation^19, 21^. In addition, transcriptional profiles for *phoA* also suggests the function of this enzyme may have also diversified, unlike *phoX.* In agreement, the transcriptomic and proteomic data in our previous study^24^ and experimental data presented here show no essential role for either *F. johnsoniae phoA* homolog in P scavenging. Given *pafA* and *phoA* can also mineralise phosphodiesters and phosphotriesters^17, 18^, these enzymes may play a large and previously unrecognised role in environmental P cycling.

*Bacteroidetes* are major organic polymer degraders that are typically associated with algal, plant and animal related niches and their success in diverse environments is driven through their ability to coexist through divergent C acquisition strategies^26, 44^. Our data reveals another unique strategy for scavenging organic C from various phosphorylated molecules that are abundant in nature^5, 45^. In ocean and plant microbiomes, this metabolism may provide a competitive advantage for C-acquisition when residual Pi levels inhibit enzyme activity rendering these molecules inaccessible. In animals, nutritional utilisation of phosphorylated carbohydrates and other organic P substrates, as well as simple sugars, plays a significant role in various pathogen-host interactions with uptake of these nutrients typically requiring the presence of specialised transporters^46–49^. This often precludes the requirement for extracellular phosphatases that are otherwise not expressed or inhibited by exogenous Pi^14, 19, 20, 22, 23^. *Bacteroidetes* are unique among bacteria lacking most ATP-binding cassette (ABC) transporters required for the uptake of organic molecules. Therefore, we speculate PafA may have evolved early in this lineage to compensate for their lack of ABC transport systems which would explain the ubiquitous occurrence of this enzyme across this phylum and why it is not associated with any particular environmental niche, unlike other *Bacteroidetes* PMEs^24^. Importantly, preferential use of phosphorylated carbohydrates as C and energy sources and subsequent release of mineralised Pi may drive P flux in systems dominated by *Bacteroidetes* and their role in environmental P cycling maybe comparable to their role in C cycling^4, 26, 31^.

Various PMEs are commercially utilised in agriculture to improve the nutritional value of animal grain feed and interest in their application as enzymes in releasing bioavailable Pi in soils in growing^50, 51^. A major limitation of PMEs is either their substrate specificity and/or their inhibition by exogenous Pi. The evidence that *Flavobacterium* PafA possesses extraordinarily high activity towards both artificial and natural organic P substrates and is easily expressed in a heterologous host is significant. PafA presents an exciting enzyme for biotechnological application, particularly for improving the nutritional value of animal and plant feed, as well as avenues aimed at developing sustainable agriculture and reducing our reliance on unsustainable chemical P fertilisers^1, 6^. Given other plant associated *Flavobacterium* spp. also express comparable or greater PME activity in the presence of exogenous Pi^24^, it is likely that PafA in these strains also possesses high activity. Given all homologs share common key residues, further investigation should ascertain the structure-function relationships responsible for the apparent differences in PME activity between various PafA homologs investigated here with the aim of enhancing their activity for commercial application.

In summary, this study resolved the contribution of seemingly redundant PMEs towards growth on organic P substrates as sole C, P and energy sources in plant associated *Flavobacterium* spp. The emergence of PafA as a highly active Pi-insensitive enzyme facilitating the rapid mineralisation of bio-available Pi that is widespread in nature signifies a major enzyme in the global P cycle.

## Materials and Methods

### Growth and maintenance of bacterial strains

*Flavobacterium johnsoniae* DSM2064 (UW101) was purchased from the Deutsche Sammlung von Mikroorganismen und Zellkulturen (DSMZ). The *Pseudomonas putida* BIRD-1 *phoX* knockout mutant was previously generated in^34^. The marine *Bacteroidetes* spp., *Algoriphagus machipongonensis* PR1 (DSM24695) and *Formosa agariphila* KMM 3901 (DSM15362) were purchased from the DSMZ collection, whereas *Polaribacter* sp. MED152 and *Gramella forsetii* KT0803 were kindly obtained from Prof Pinhassi and Dr Wulf, respectively. The Roseobacter strains were historically obtained from the DSMZ collection. *F. johnsoniae* and *P. putida* genotypes were routinely maintained on casitone yeast extract medium (CYE)^52^ containing casitone (4 g ^L-1^), yeast extract (1.25 g ^L-1^), MgCl2 (350 mg ^L-1^) and 20 g ^L-1^ agar or lysogeny broth (LB) containing 15 g L^-1^ agar, respectively. For various growth experiments and phosphatase assays, *F. johnsoniae* was grown in a minimal A medium adapted from^19^. This medium contained glucose 5-20 mM, NaCl 200 mg ^L−1^, NH4Cl 450 mg ^L−1^, CaCl2 200 mg ^L−1^, KCL mg ^L−1^ MgCl2 450 mg ^L−1^, FeCl2 10 mg ^L−1^, MnCl2 10 mg ^L−1^, 20mM Bis/Tris buffer pH 7.2. KH2PO4 added to a final concentration ranging from 50 μM to 1 mM. For *P. putida* genotypes, glucose was replaced with sodium succinate (final concentration 15 mM). Marine *Bacteroidetes and* Roseobacter strains were grown in Difco Marine Broth 2216 (Fischer Scientific) and incubated at 28°C.

The organic P substrates fructose 6-phosphate, glucose 6-phopshate, glycerol phosphate, sodium phosphorylcholine, sn-Glycerol 3-phosphate bis(cyclohexylammonium), phosphocholine chloride calcium salt tetrahydrate and L-α-Phosphatidylinositol (∼50% TLC) from Glycine max (soybean) ∼50% (TLC) were also purchased from Sigma-Aldrich, Merck. For various growth experiments using organic P substrates as a sole P source 200-500 μM was added to the minimal medium. For growth experiments using organic P substrates as a sole C source or sole C and P source, 2-3 mM was added to the minimal medium.

For Pi mineralisation experiments generating conditioned medium, glucose (5 mM) and either 1 mM glucose 6-phopshate or 100 μM Pi was used. After overnight growth, supernatants were collected after removing cells by centrifugation (10,000 x *g*) and filtration through a PES membrane (0.22 μm pore size). Spent medium was mixed with fresh medium containing 5 mM glucose (50:50% v/v) and half of the subsequent culture lines were supplemented with 250 μM Pi (positive control).

### Construction of *Flavobacterium* mutants

To construct the various PME mutants in DSM2064, the method developed by^53^ was used. A full list of primers used in this study can be found in Table S14. Briefly, two 1-1.6 kb regions flanking each gene were cloned into plasmid pYT313 using the HiFi assembly Kit (New England Biosciences). Sequence integrity was checked via sequencing. The resulting plasmids were transformed into the donor strain *Escherichia coli λ*S17-1 (S17-1) and mobilised into *Flavobacterium* via conjugation (overnight at 30°C). 5 ml overnight cultures were used to inoculate (25% v/v) fresh 5 ml CYE or LB media and incubated for 8 h. A 200 μL donor: recipient suspension (1:1) was plated onto CYE containing erythromycin (100 μg mL^-1^). Colonies were re-streaked onto CYE erythromycin plates to remove any background wild type. Single homologous recombination events were confirmed by PCR prior to overnight growth in CYE followed by plating onto CYE containing 10% (w/v) sucrose to select for a second recombination event. To identify a double homologous recombination mutant, colonies were re-grown on CYE containing 10% (w/v) sucrose and CYE containing erythromycin 100 μg mL^-1^. Erythromycin sensitive colonies were screened for a double homologous recombination event by PCR targeting a region deleted in the mutant.

### Construction of *Pseudomonas putida* BIRD-1 strains

To complement the *Pseudomonas* sp. BIRD1 *ΔphoX* mutant with various *Bacteroidetes* phosphatases, the promoter for the native *phoX*:BIRD-1, was cloned into a broad-host range plasmid, pBBR1MCS-km, using methods outlined in^34^. The various *Flavobacterium* PMEs were subsequently cloned downstream of this promoter using the HIFI assembly kit (New England Biosciences). For PafA homologs found in *Chitinophaga pinensis* DSM2588 (CP1, 644962876; CP2, 644963845), the open reading frame of each gene was chemically synthesised (Integrated DNA Technologies, gBlocks gene fragment service) with *Hin*dIII and *Xba*I restriction sites added at the 5’ and 3’ ends, respectively. IMG gene accession numbers are given in parentheses. After digestion of fragment and plasmid, ligation was performed using T4 DNA ligase (Promega). Plasmids were mobilised into the *ΔphoX* mutant via electroporation using a voltage of 18 kv cm^-1^ or by bi-parental mating with the donor strain S17-1. For electroporation, cells were immediately added to LB and incubated for 2-3 h prior to selection on LB supplemented with 50 μg mL^-1^ kanamycin. For conjugative plasmid transfer, S17-1 and *P.putida* (*ΔphoX::Gm*) were grown overnight in LB broth (0.5 ml) and resuspended in 0.1 ml fresh medium. Strains were mixed and spotted onto LB agar and incubated for 5 h. Cells were scraped from solid medium and resuspended in 1 ml Tris HCL buffer (pH 7.4). A 1:10 serial dilution was established and plated onto LB supplemented with 50 μg mL^-1^ kanamycin and 10 μg mL^-1^ gentamicin to counter select against S17-1. Colonies were screened by PCR for the presence of the plasmid.

### Quantification of alkaline phosphatase activity

The protocol was adapted from^24^, where volumes were adjusted for compatibility with a microtiter plate reader (Tecan SPARK 10M). Cell cultures (n = 3) for both Pi-replete and Pi-deplete growth conditions were directly incubated (30°C at 160 rpm) with 10 mM (final. concentration) *para*-nitrophenyl phosphate (pNPP) or resuspended in a Tris-HCl buffer adjusted to pH 5.4, 7.4 or 9.4 prior to pNPP incubations. All reactions were incubated at 28°C on a rotary shaker (230 rpm). The reaction was stopped using 2 mM (final concentration) NaOH once a colour change was detected, and reactions were still operating in a linear manner. Reactions were spent when absorbance (405nm) reached >4 and all reactions were stopped when 5-25% of substrate was consumed, typically within 2–60 min. For each strain and growth condition, A405nm measurements were corrected by subtracting A405nm measurements for reactions immediately stopped with NaOH. Normalisation against the culture optical density (OD600) was performed and the rate was calculated and expressed _h_−1.

### Quantification of exogenous phosphate

To quantify Pi mineralisation in the presence of organic P, cells were grown in a minimal medium supplemented with glucose (5 mM) and 2 mM of either glycerol phosphate and sodium phosphorylcholine (50:50% mix) or fructose 6-phosphate and glucose 6-phopshate (50:50% mix). A control using 500 μM Pi was also established. To quantify exogenous Pi, cells were removed from culture aliquots via centrifugation (10000 x *g* for 5 min), and Pi concentrations in the supernatants were determined according to the method of Chen *et al*.^54^ and further modified here for compatibility with a microtiter plate reader. Briefly, 100 μL supernatant was added to 100 μL of dH2O:6N sulfuric acid:2.5% w/v ammonium molybdate:10% w/v ascorbic acid (2:1:1:1). For each individual assay, absolute quantification of Pi was achieved using a standard curve of known Pi concentrations (0, 7.8125, 15.625, 31.25, 62.5, 125, 250, 500 μM). This enabled a reduced incubation at 37°C (30-45 min). Absorbance was measured at 820nm. Each *F. johnsoniae* strain, wild type plus three PME mutants, were grown under the different treatments in duplicate cultures with duplicate technical replicates for growth and Pi concentrations taken for each culture (n=4 total).

### Bioinformatics analyses

The online platform Integrated Microbial Genomes & Microbiomes server at the Joint Genome Institute (IMG/JGI) was used to perform most comparative genomics analyses described in this study. Genomes and metagenomes were stored in Genome sets and for PafA, BLASTP searches (Min. similarity 30%, E-value e^−50^) were set up using the ‘jobs function’. The diversity, richness, gene and transcript abundance of each phosphatase, PhoA, PhoD, PhoX and PafA in seawater was determined by searching the TARA ocean metagenome (OM-RGC_v2_metaG) and metatranscriptome (OM-RGC_v2_metaT) via the Ocean Gene Atlas web interface, using the hmmsearch function (stringency 1E^-60^). profile Hidden Markov Models (pHMM) for PhoA (PF00245), PhoX (PF05787) and PhoD (PF PF09423), were downloaded from https://pfam.xfam.org/. For PafA, a pHMM was manually curated by aligning sequences using MUSCLE identified in various *Bacteroidetes* isolates and pHMM using the hmmbuild function in hmmer 3.3 (http://hmmer.org). Sequence abundances were expressed as the average percentage of genomes containing a gene copy or transcript by dividing the percentage of total mapped reads by the median abundance (as a percentage of total mapped reads) of 10 single-copy marker genes^55^ for both MG and MT.

To determine the phylogeny of PafA, sequences were aligned using MUSCLE and manually inspected for the possession of key residues using MEGAX. Sequences possessing any point mutations were removed from the multiple alignment. Phylogenetic reconstruction was performed using IQ-Tree using the parameters -m TEST -bb 1000 -alrt 1000. Evolutionary distances were inferred using maximum-likelihood analysis. Relationships were visualised using the online platform the Interactive Tree of Life viewer (https://itol.embl.de/).

All statistical analyses and data visualisation were performed using the ggplot2, ggfortify, tidyr, plyr, serration, rcolourbrewer packages in Rstudio (1.2.5033).

## Acknowledgements

This study was funded by the Biotechnology and Biological Sciences Research Council (BBSRC) under project code BB/T009152/1 linked to a Discovery Fellowship (IL) and a Rank Prize Fund New Lecturer Award.

## Supplementary Information

### Supplementary Tables

**Table S1.**
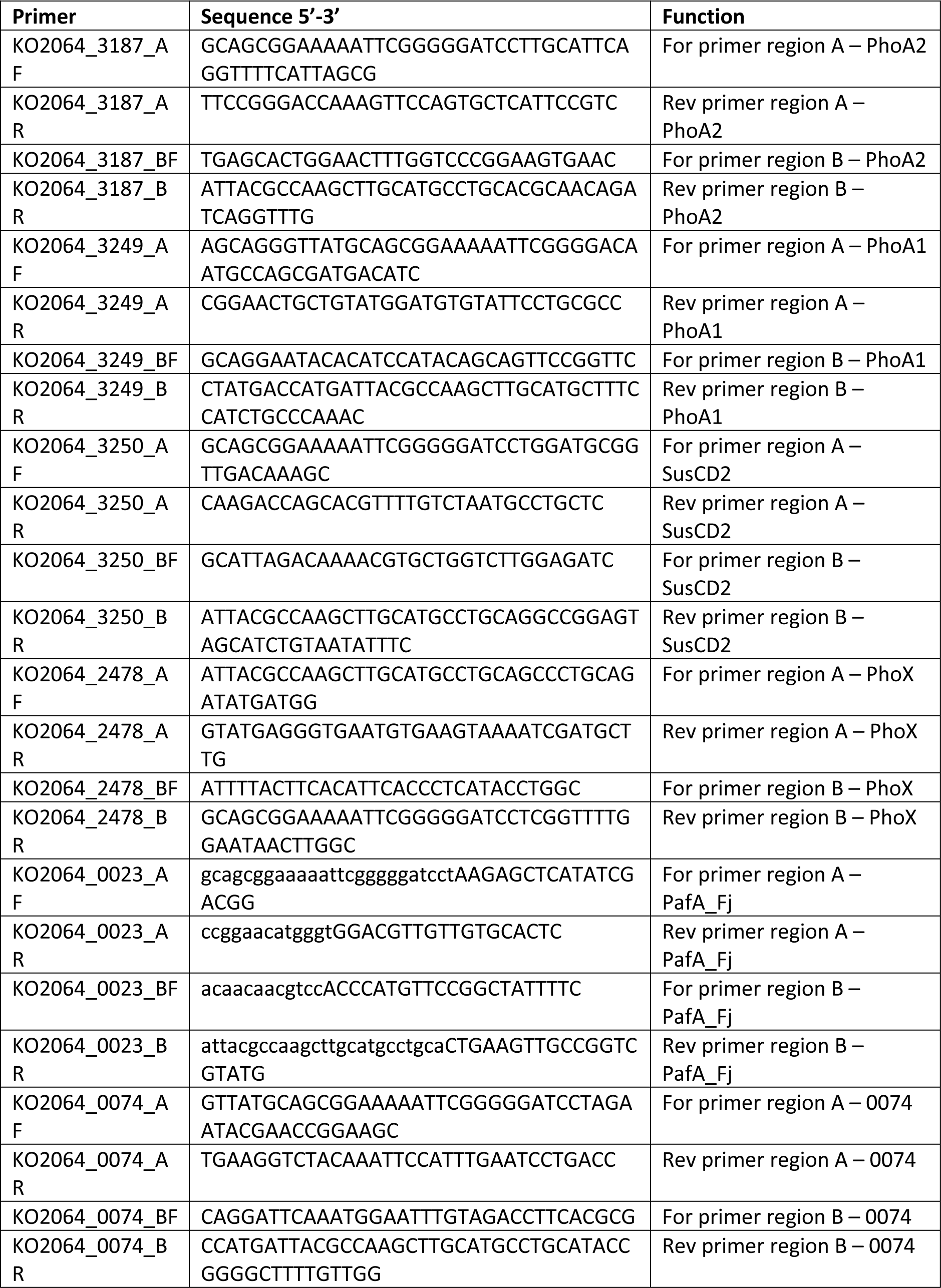
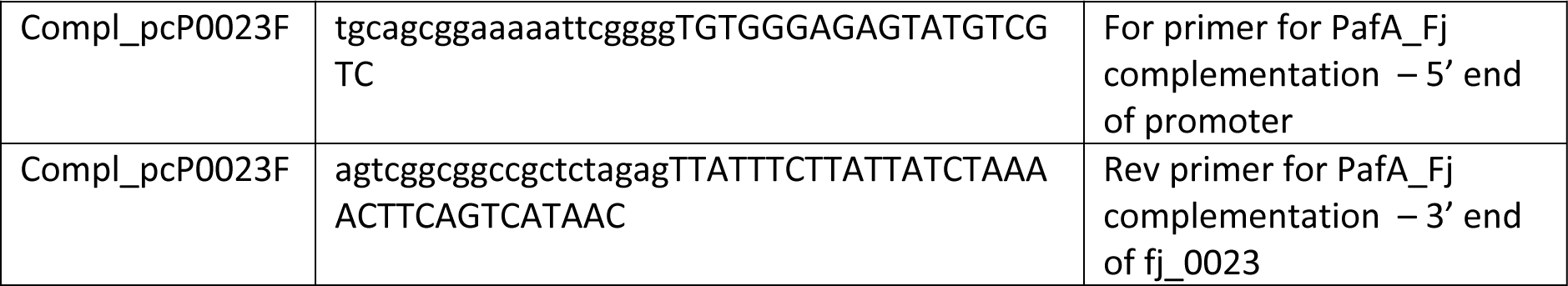
List of primers used for mutagenesis of *Flavobacterium johnsoniae* DSM2064

**Table S2.** List of strains used and generated in this study:

| Strain | Description *fjoh_0023* | Reference |
| --- | --- | --- |
| *F. johnsoniae* DSM2064 | Wild type strain | DSMZ |
| *P. putida* BIRD-1 | Wild type strain | (3) |
| *Polaribacter* sp. MED152 | Wild type marine *Flavobacteriia* (*Bacteroidetes*) | (4) |
| *Gramella forsetii* KT0803 | Wild type marine *Flavobacteriia* (*Bacteroidetes*) | (5) |
| *Formosa agariphila* KMM 3901 | Wild type marine *Flavobacteriia* (*Bacteroidetes*) | (6) |
| *Algoriphagus machipongonensis* PR1 | Wild type marine *Cytophagia* (*Bacteroidetes*) | (7) |
| *Ruegeria pomeroyi* DSS-3 | Wild type marine *Rhodobacteraceae* (*Alphaproteobacteria*) | DSMZ |
| *Dinoroseobacter shibae* DFL-12 | Wild type marine *Rhodobacteraceae* (*Alphaproteobacteria*) | DSMZ |
| *Roseobacter denitrificans* OCh 114 | Wild type marine *Rhodobacteraceae* (*Alphaproteobacteria*) | DSMZ |
| $\Delta phoX::Gm$ | Mutant strain fo *P. putida* with *PPUBIRD1_1093 phoX* mutated | (8) |
| $\Delta phoX::Gm$ +pBX:*phoX*<sup>BIRD</sup> | $\Delta phoX::Gm$ complemented with pBBR1MCS-km containing the *phoX* promoter (+pBX) and its native *phoX* | (8) |
| $\Delta phoX::Gm$ +pBX:*phoA*<sup>Ec</sup> | $\Delta phoX::Gm$ complemented with +pBX containing the *E. coli phoA* | This study |
| $\Delta phoX::Gm$ +pBX:*phoA1*<sup>Fj</sup> | $\Delta phoX::Gm$ complemented with +pBX containing *fjoh_3249 (phoA1)* | This study |
| $\Delta phoX::Gm$ +pBX:*phoA2*<sup>Fj</sup> | $\Delta phoX::Gm$ complemented with +pBX containing *fjoh_3187 (phoA2)* | This study |
| $\Delta phoX::Gm$ +pBX:*phoX*<sup>Fj</sup> | $\Delta phoX::Gm$ complemented with +pBX containing *fjoh_2478 (phoX)* | This study |
| $\Delta phoX::Gm$ +pBX:*pafA*<sup>Fj</sup> | $\Delta phoX::Gm$ complemented with +pBX containing *fjoh_0023 (pafA)* | This study |
| *phoX::Gm* +pBX:*pafA*<sup>Fj</sup> | $\Delta phoX::Gm$ complemented with +pBX containing *Cpin_0724 (pafA1)* | This study |
| $\Delta phoX::Gm$ +pBX:*pafA*<sup>Fj</sup> | $\Delta phoX::Gm$ complemented with +pBX containing *Cpin_1665 (pafA2)* | This study |
| $\Delta pafA$ | *F. johnsoniae* with *fjoh_0023 (pafA)* mutated | This study |
| $\Delta phoX^{Fj}$ | *F. johnsoniae* with *fjoh_2478 (pafA)* mutated | This study |
| $\Delta 0074$ | *F. johnsoniae* with *fjoh_0074 (pafA)* mutated | This study |
| $\Delta phoA1:\Delta phoA2$ | *F. johnsoniae* with *fjoh_3187 (phoA2)* and *fjoh_3249 (phoA2)* mutated | This study |
| Quad ($\Delta phoA1:\Delta phoA2:\Delta phoX^{Fj}:\Delta 0074$) | *F. johnsoniae* with *fjoh_0074*, *fjoh_2478*, *fjoh_3187 (phoA2)* and *fjoh_3249 (phoA2)* mutated | This study |
| M5 ($\Delta phoA1:\Delta phoA2:\Delta phoX^{Fj}:\Delta 0074:\Delta pafA$) | *F. johnsoniae* with *fjoh_0023*, *fjoh_0074*, *fjoh_2478*, *fjoh_3187 (phoA2)* and *fjoh_3249 (phoA2)* mutated | This study |
| M5 +pCP:*pafA* | M5 +pCP11 containing *fjoh_0023* and its native promoter | This study |

**Table S3.** Search-algorithm comparison of *phoX* and *phoA* gene abundance in the TARA Oceans dataset. For BLASTP searches, the same query sequences and search parameters as Sebastien et al. (2009) were used.

| Phosphatase | No. of hits | No. of hits | Abundance value | Abundance value |
| --- | --- | --- | --- | --- |
|  | BLASTP | hmmer | BLASTP | hmmer |
| *phoA* | 51 | 947 | 1020 | 26114 |
| *phoX* | 409 | 687 | 8951 | 17135 |

All searches were performed using a stringency $e^{-60}$ .

### Supplementary Figures

**Figure S1.**
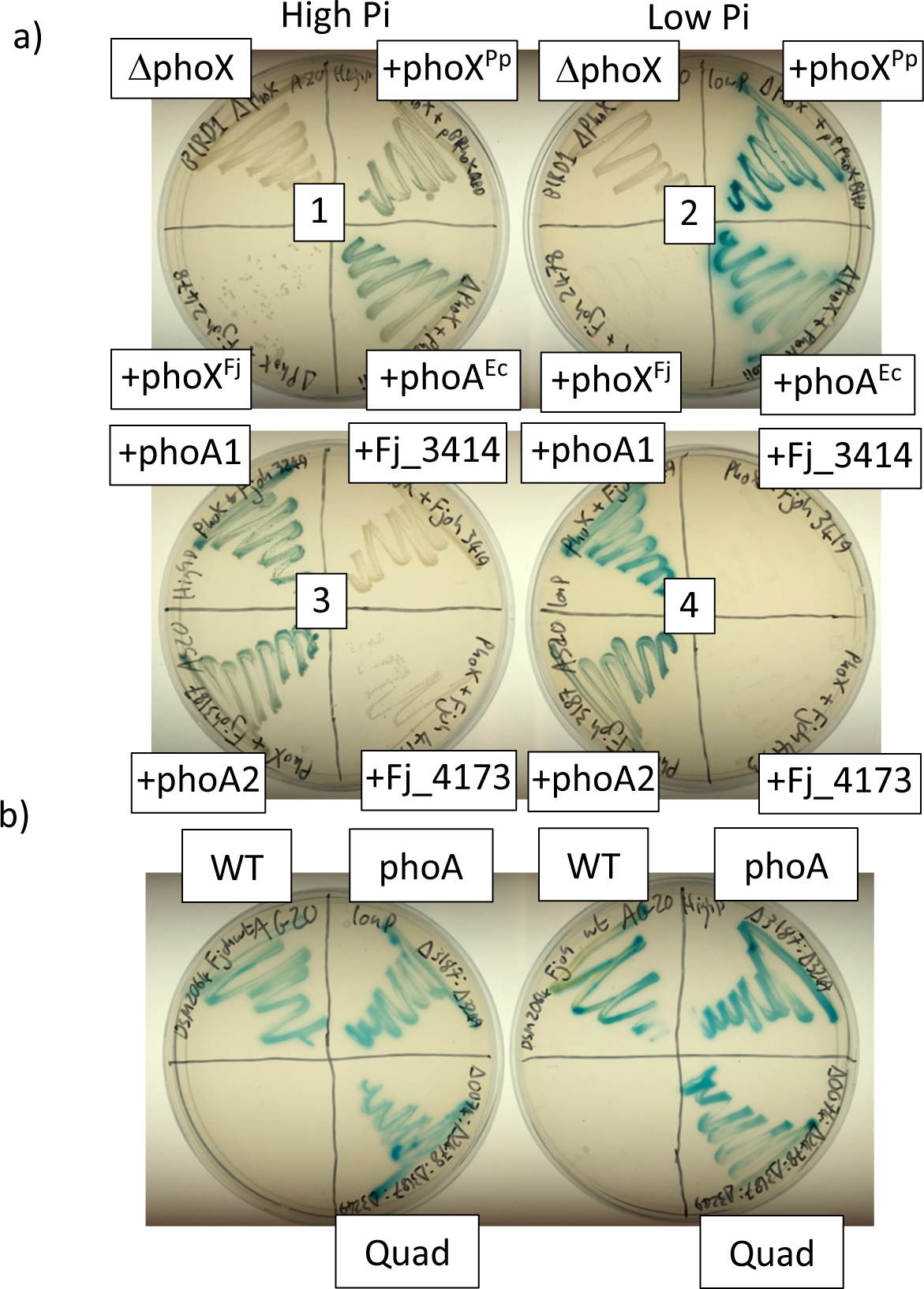
Qualitative plate assay for phosphomonoesterase acitvity. (a) Alkaline phosphatase plate assay using the *Pseudomonas putida* BIRD-1 *ΔphoX* mutant (top left, plates 1 & 2). 5-Bromo-4-chloro- 3-indolyl phosphate (BCIP), more commonly known as XP, was used as the substrate. The *phoX* mutant exhibited zero activity indicated by a lack of blue colour which is created when XP is cleaved. Complementation with the native *P. putida* BIRD-1 (+phoX^Pp^) restored the wild type phenotype. Heterologous expression of the two PhoA-like homologs (Fjoh_3187, bottom left P3 & 4 and Fjoh_3249, top left P3 & 4), also restored APase activity confirming their function. Neither Fjoh_3414 nor Fjoh_4173 restored any phenotype. Interestingly, Fjoh_2478, the PhoX-like homolog, did not restore the phenotype. However, growth in this complemented mutant was inhibited which suggests that expression and subsequent export of the lipoprotein may be affected. Plates were left overnight at 30°C. **(b)** Alkaline phosphatase plate assay using the various *F. johnsoniae* APase mutants. Both the double *phoA* mutant and the quadruple mutant still displayed observable APase activity under Pi replete and Pi deplete growth conditions (top left, plates 1 & 2). 5-Bromo-4-chloro-3-indolyl phosphate (BCIP), more commonly known as XP, was used as the substrate. Abbreviations: WT, wild type; *phoA*, *ΔphoA1: ΔphoA2;* Quad, *ΔphoA1:ΔphoA2:ΔphoX:Δfjoh_0074*.

**Figure S2.**
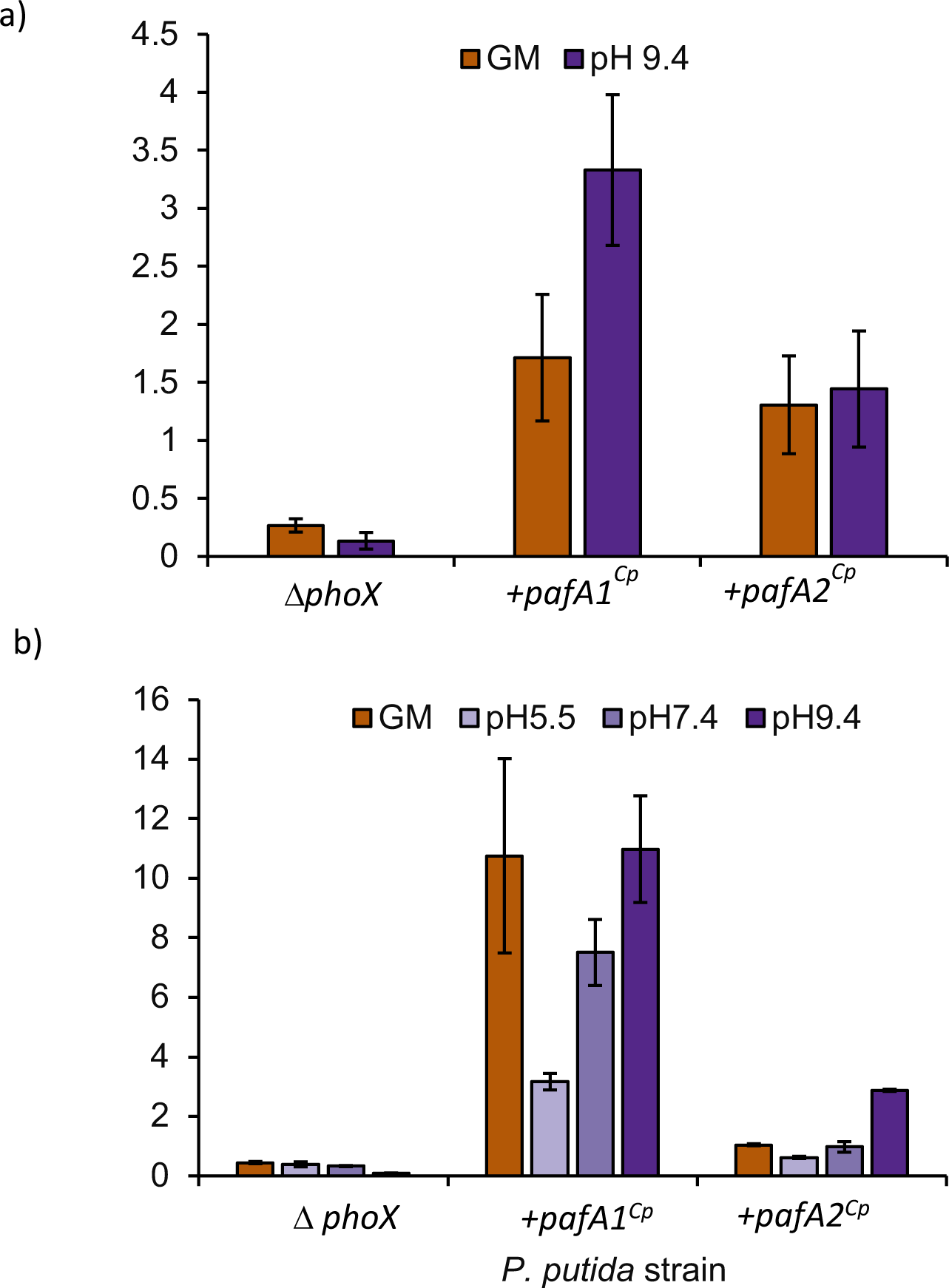
Heterologous phosphomonoesterase (PME) activity in the *P. putida* null mutant. Cell expressing two distinct *pafA* homologs found in the genome of *Chitinophaga pinensis* (+*pafA1^Cp^* +*pafA2^Cp^*) were grown overnight in complex medium (**a**) or minimal medium **(b)** established phosphate-deplete (Low Pi) growth conditions. PME activity was obtained through addition of the artificial substrate *para*-nitrophenyl phosphate (10mM). Values presented are the mean of biological triplicates and error bars denote standard deviation. Abbreviations: GM, growth medium

**Figure S3.**
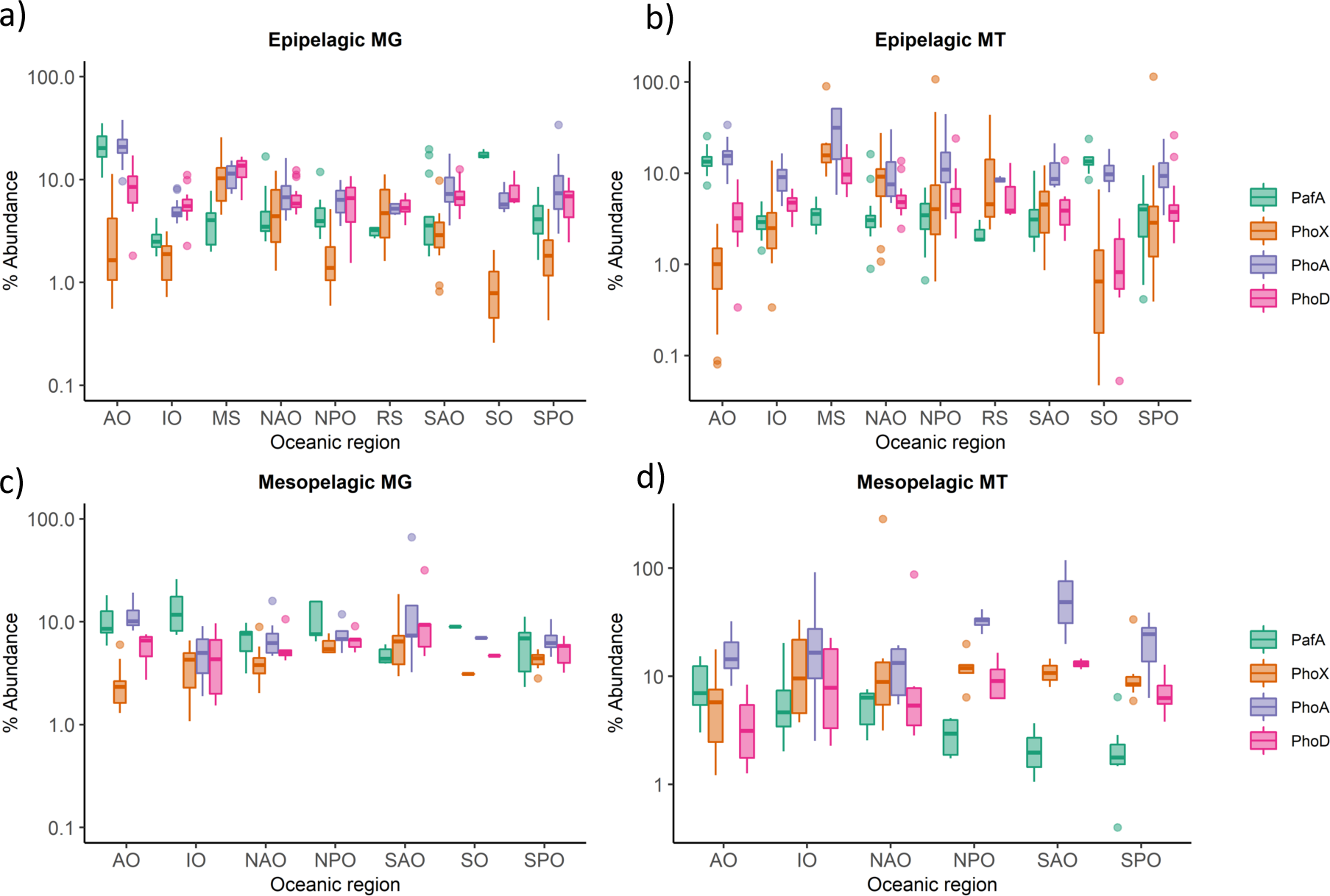
Distribution and expression of phosphatase genes across specific regions of the global ocean. Abundance (Log10 abundance [gene or transcript] relative to the median abundance [gene or transcript] of 10 single copy core genes) of *pafA*, *phoX*, phoA, *phoD* in marine epipelagic (**a, b**) and mesopelagic (**c, d**) waters, split by metagenome (MG) (**a, c**) and metatranscriptome (**b, d**). Data are represented as boxplots, where the middle line is the median and the upper and lower hinges correspond to the first and third quartiles. The upper whisker extends from the upper hinge to the largest value that is no more than 1.5×IQR (inter-quartile range) from the upper hinge, and the lower whisker extends from the lower hinge to the smallest value that is no further than 1.5×IQR from the lower hinge. Data beyond the ends of the whiskers are outlier points that are plotted individually.

**Figure S4.**
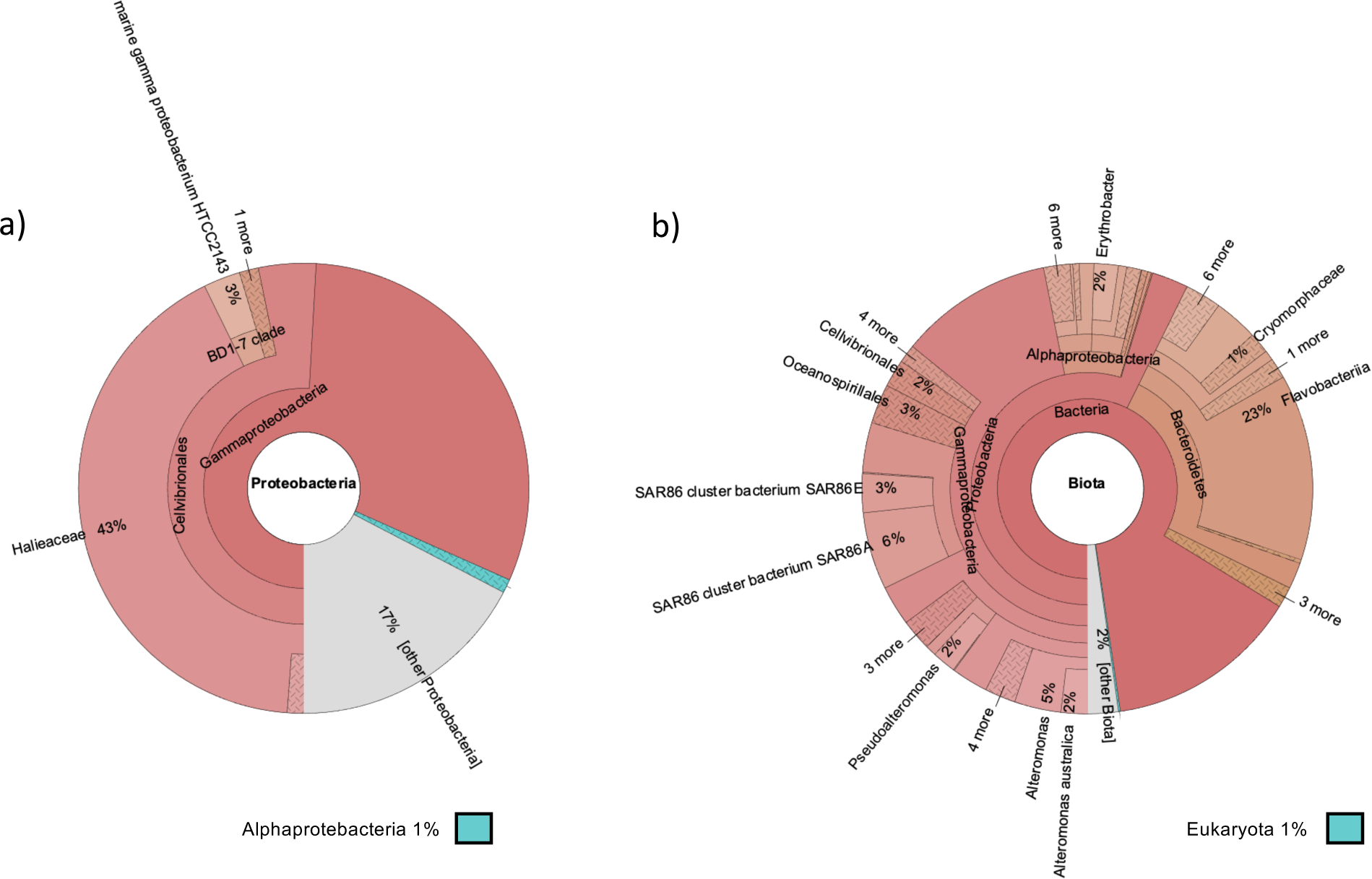
Taxonomic classification of PhoA homologs retrieved from the global ocean. The TARA Oceans OM-RGCv2+G metagenome dataset was scrutinised using either BLASTP (**a**) or hmmer (**b**) search algorithms. Both stringency values were set at e^-60^.

**Figure S5.**
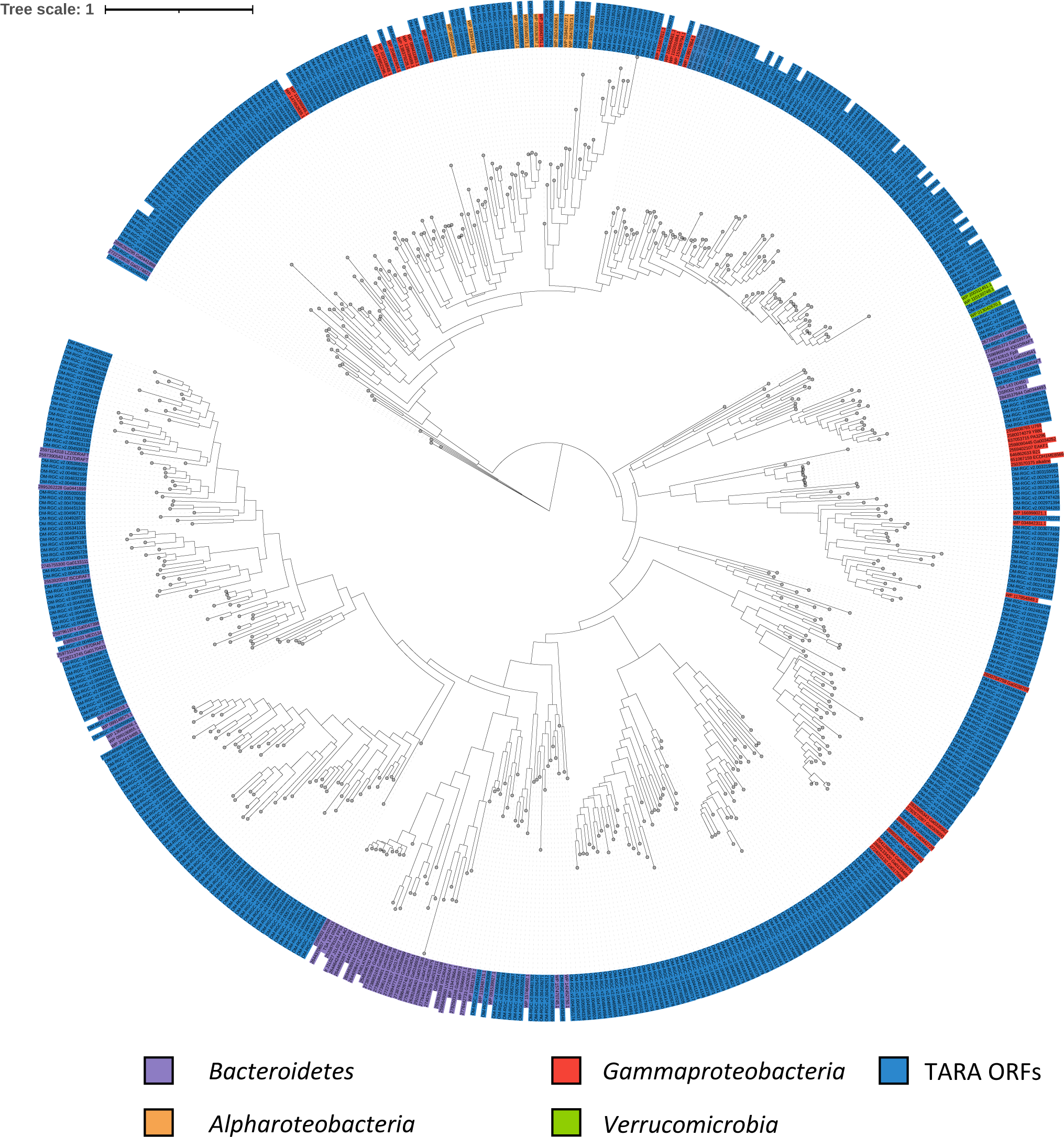
Phylogeny of PhoA homologs retrieved from the TARA oceans dataset. Tree topology and branch lengths were calculated by maximum likelihood using the Blosum62+F+G4 model of evolution for amino acid sequences based on 900 sites in lQ-TREE software. A consensus tree was generated using 1000 bootstraps. TARA ORFs are coloured navy blue. Sequences retrieved from isolate genomes (IMG gene numbers or NCBI 11accessions provided) were also included and colours represent taxonomy (see legend).

**Figure S6.**
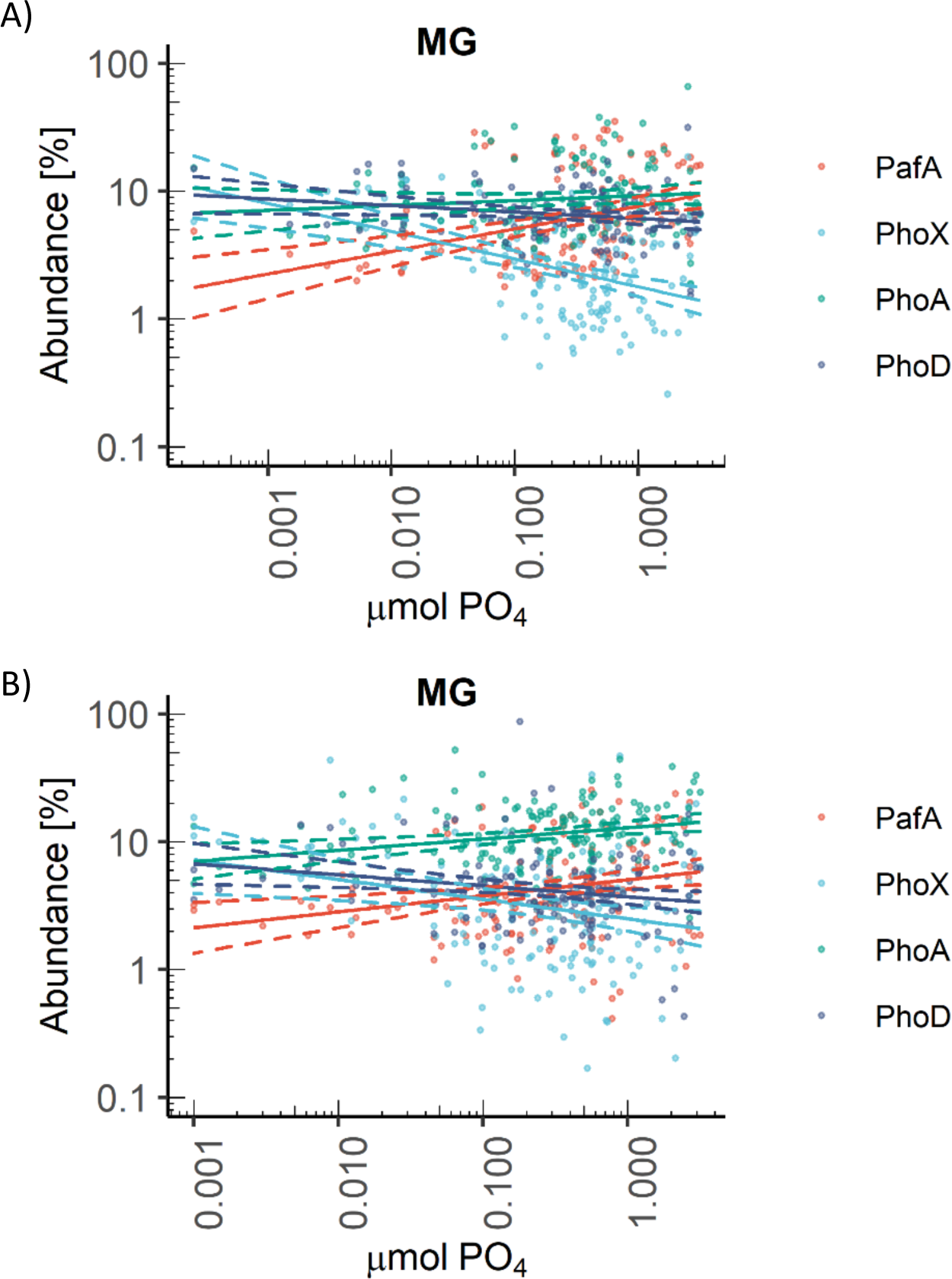
Relationship between gene and transcript abundance of *pafA*, *phoX, phoA*, *phoD* and standing stock phosphate concentrations in the global ocean. Phosphatase abundance, analysed by linear regression of Log10 gene abundance and standing stock phosphate (PO4) concentrations, in the epipelagic MG (**a**) and mesopelagic MG **(b)**. 95% confidence intervals are shown by dashed lines.

